# Genetic assimilation of ancestral plasticity during parallel adaptation

**DOI:** 10.1101/2021.11.30.470425

**Authors:** Daniel P. Wood, Jon A. Holmberg, Owen G. Osborne, Andrew J. Helmstetter, Luke T. Dunning, Amy R. Ellison, Rhian J. Smith, Jackie Lighten, Alexander S.T. Papadopulos

## Abstract

Phenotypic plasticity in ancestral populations is hypothesised to facilitate adaptation, but evidence supporting its contribution is piecemeal and often contradictory. Further, whether ancestral plasticity increases the probability of parallel genetic and phenotypic adaptive changes has not been explored. The most general finding is that nearly all ancestral gene expression plasticity is reversed following adaptation, but this is usually examined transcriptome-wide rather than focused on the genes directly involved in adaptation. We investigated the contribution of ancestral plasticity to adaptive evolution of gene expression in two independently evolved lineages of zinc-tolerant *Silene uniflora*. We found that the general pattern of reversion is driven by the absence of a widespread stress response in zinc-adapted plants compared to ancestral, zinc-sensitive plants. Our experiments show that reinforcement of ancestral plasticity plays an influential role in the evolution of plasticity in derived populations and, surprisingly, one third of constitutive differences between ecotypes are the result of genetic assimilation of ancestral plasticity. Ancestral plasticity also increases the chance that genes are recruited repeatedly during adaptation. However, despite a high degree of convergence in gene expression levels between independently adapted lineages, genes with ancestral plasticity are as likely to have similar expression levels in adapted populations as genes without. Overall, these results demonstrate that ancestral plasticity does play an important role in adaptive parallel evolution, particularly via genetic assimilation across evolutionary replicates.

## Introduction

The contributions of determinism and contingency in shaping evolution are hotly debated (Gould, 1989; Conway Morris, 2003; Orgogozo, 2015). Whether repeated adaptation to the same environment results in similar changes at the molecular level is key to understanding this balance (Gould, 1989; Christin *et al*., 2010; Losos, 2011; Bolnick *et al*., 2018), as well as the predictability of future responses to environmental change (Waldvogel *et al*., 2020). Adaptation to novel environments often involves gene expression changes, but previous studies have found varying degrees of parallelism during repeated adaptation (Hanson *et al*., 2017; Stern & Crandall, 2018; Parker *et al*., 2019; Jacobs *et al*., 2020). These changes occur at various levels, including in the overlap of shared differentially expressed genes, fold-changes of these genes, or final expression levels (Ghalambor *et al*., 2015; Jacobs *et al*., 2020). Understanding the mechanisms that influence the extent of parallelism is an important step in predicting evolutionary responses to new environmental challenges (Stern, 2013; Bolnick *et al*., 2018; Waldvogel *et al*., 2020).

Phenotypic plasticity in ancestral populations (i.e., ancestral plasticity) is suspected to play a role in facilitating adaptation to new environments (Baldwin, 1896; Ghalambor *et al*., 2007; Schaum *et al*., 2013). In addition to generally preserving the genetic variability of a colonising population (Draghi & Whitlock, 2012), plastic responses to new environments could provide the basis for adaptation by moving the trait values in some individuals closer to the new local optimum (Levis *et al*., 2018). Beneficial plasticity of this kind could be retained in locally adapted populations or genetically assimilated and canalised into constitutive expression differences (Heckel *et al*., 2016). Alternatively, ancestral plasticity that takes expression levels further away from the new optimum is potentially maladaptive and could hinder adaptation to the novel environment (Velotta *et al*., 2018; Josephs *et al*., 2021).

Current evidence suggests a variety of possible impacts of ancestral plasticity on adaptation (Ghalambor *et al*., 2015; Kenkel & Matz, 2017; Ho & Zhang, 2018; Velotta *et al*., 2018; Kelly, 2019), but the relationship between plasticity and evolutionary parallelism has received limited attention (Oke *et al*., 2016; Bolnick *et al*., 2018). Other properties of gene expression in ancestral populations, such as ancestral expression level or tissue expression location, are associated with increased co-option and potentially parallelism (Hargreaves *et al*., 2014; Moreno-Villena *et al*., 2018). If phenotypic plasticity significantly facilitates the repurposing of traits during adaptation (Moczek *et al*., 2011), then beneficial plasticity may result in greater parallelism than when plasticity is maladaptive.

Previous studies have generally found that most ancestral plasticity across transcriptomes is reversed in derived populations, taking expression values further from the new optimum (Ho & Zhang, 2018, 2019; Swaegers *et al*., 2020; Fischer *et al*., 2021; although see Mäkinen *et al*., 2016; Mallard *et al*., 2020). However, there are examples of ancestral plasticity in particular genes or traits facilitating subsequent adaptation (Scoville & Pfrender, 2010; Levis *et al*., 2018; Velotta *et al*., 2018; Wang & Althoff, 2019). Most expression studies on the topic examine transcriptome-wide patterns in ancestrally plastic genes, rarely considering whether genes involved in evolutionary adaptation to the new environment are more likely to have possessed beneficial ancestral plasticity, when compared to the whole transcriptome (Ho & Zhang, 2018; Koch & Guillaume, 2020; Swaegers *et al*., 2020; Josephs *et al*., 2021; Fischer *et al*., 2021; Bittner *et al*., 2021). Transcriptome-wide assessments include changes that may not directly contribute to adaptation (in the evolutionary sense), such as those stemming from general stress responses. As a result, estimates of the contribution of ancestral plasticity to adaptation may be distorted in whole transcriptome analysis.

Here, we investigate the relationship between ancestral plasticity, adaptation and parallelism using independently evolved lineages of zinc-tolerant *Silene uniflora* from contaminated metal mines and local zinc-sensitive coastal populations (Papadopulos *et al*., 2021). In this species, ancestral coastal populations have repeatedly colonised contaminated mine soils throughout Great Britain and Ireland over the past 250 years (Baker, 1974), producing locally adapted populations that can grow at high concentrations of zinc (Baker, 1974, 1978; Papadopulos *et al*., 2021). This provides an ideal opportunity to investigate the role of ancestral plasticity in adaptation across multiple evolutionary replicates.

## Results and Discussion

We quantified gene expression in the roots of two independently derived, zinc-tolerant populations from geographically distant, derelict mines (T1 - England, T2 - Wales) and their nearest and most closely related zinc-sensitive coastal populations which represent the ancestral phenotype (S1 and S2; Fig. S1). Extant zinc-sensitive coastal populations were used as proxies for ancestral expression, which was compared to responses in recently adapted zinc-tolerant mine populations. We exposed clones of the same individuals to two treatment conditions (control or zinc-contaminated) and collected RNA-seq data from the roots of the experimental plants. Our experimental design allowed us to quantify: (i) the ancestral plastic response to zinc contamination, ii) the extent of convergent gene expression changes during rapid parallel adaptation; the evolutionary response to ancestral plasticity at iii) a transcriptome-wide level and iv) for genes plausibly involved in adaptation; and v) the relationship between ancestral plasticity and convergent gene expression changes. In so doing, we establish the extent to which rapid adaptation is shaped by constraint and plasticity, disentangling the influence of general stress responses versus adaptive responses on patterns of reversion and reinforcement.

Heavy metals are highly phytotoxic and high concentrations of zinc have a considerable impact on growth and fitness of coastal populations of *S. uniflora* (Baker, 1978; Papadopulos *et al*., 2021). Transcriptome-wide ancestral plasticity (i.e., the response to zinc in sensitive populations) was dominated by a general and widespread stress response. In total, 48.0% of the transcriptome (27,607 genes) was differentially expressed in both sensitive populations between treatments, with an overwhelming majority shared across populations (Figure S2A). Shared upregulated genes were enriched for 18 GO terms related to stress (Table S1). Further, the overall major difference in expression between susceptible and tolerant populations in the zinc treatment was the lack of this stress response in tolerant populations. In total, 18,343 genes were differentially expressed in both pairs of tolerant and sensitive populations in the zinc (Figure S2B), which were enriched for 22 stress-related GO terms (Table S2). Of these genes, 83.7% were ancestrally plastic (i.e., also differentially expressed between treatments in both sensitive populations), but only 3.6% showed derived plasticity (i.e., were also differentially expressed between treatments in both tolerant populations; Figure S2C). This reveals a significant and widespread disruption to transcription in sensitive plants, consistent with the broad impact of zinc toxicity on cellular processes (Singh *et al*., 2016). It also indicates that, in general, greater transcriptomic perturbations in ancestral populations exposed to new environments may be driven by general stress responses (Koch & Guillaume, 2020; Swaegers *et al*., 2020; Josephs *et al*., 2021; Bittner *et al*., 2021).

### Rapid evolution of highly parallel gene expression changes

*Silene uniflora* has independently colonised mines and evolved tolerance to the very high levels of zinc (2,400-48,100ppm) in the contaminated soils (Baker, 1974, 1978; Papadopulos *et al*., 2021). Given that this phenotype has evolved in parallel as a result of a strong selection pressure, we also expected a component of the transcription profiles to show parallel changes in tolerant populations. In the control treatment, principal component analysis of transcriptome-wide gene expression levels revealed separation of populations by zinc-tolerance (i.e., tolerant vs. sensitive) on PC1 and by geographic origin (i.e., T1 and S1 vs. T2 and S2) on PC2 (Fig. 1A). Within-population variation was low relative to between populations/treatments. In these benign control conditions, the trajectories of whole transcriptome profile evolution were divergent and almost orthogonal rather than parallel (*sensu* Bolnick et al. 2018).

**Figure 1:**
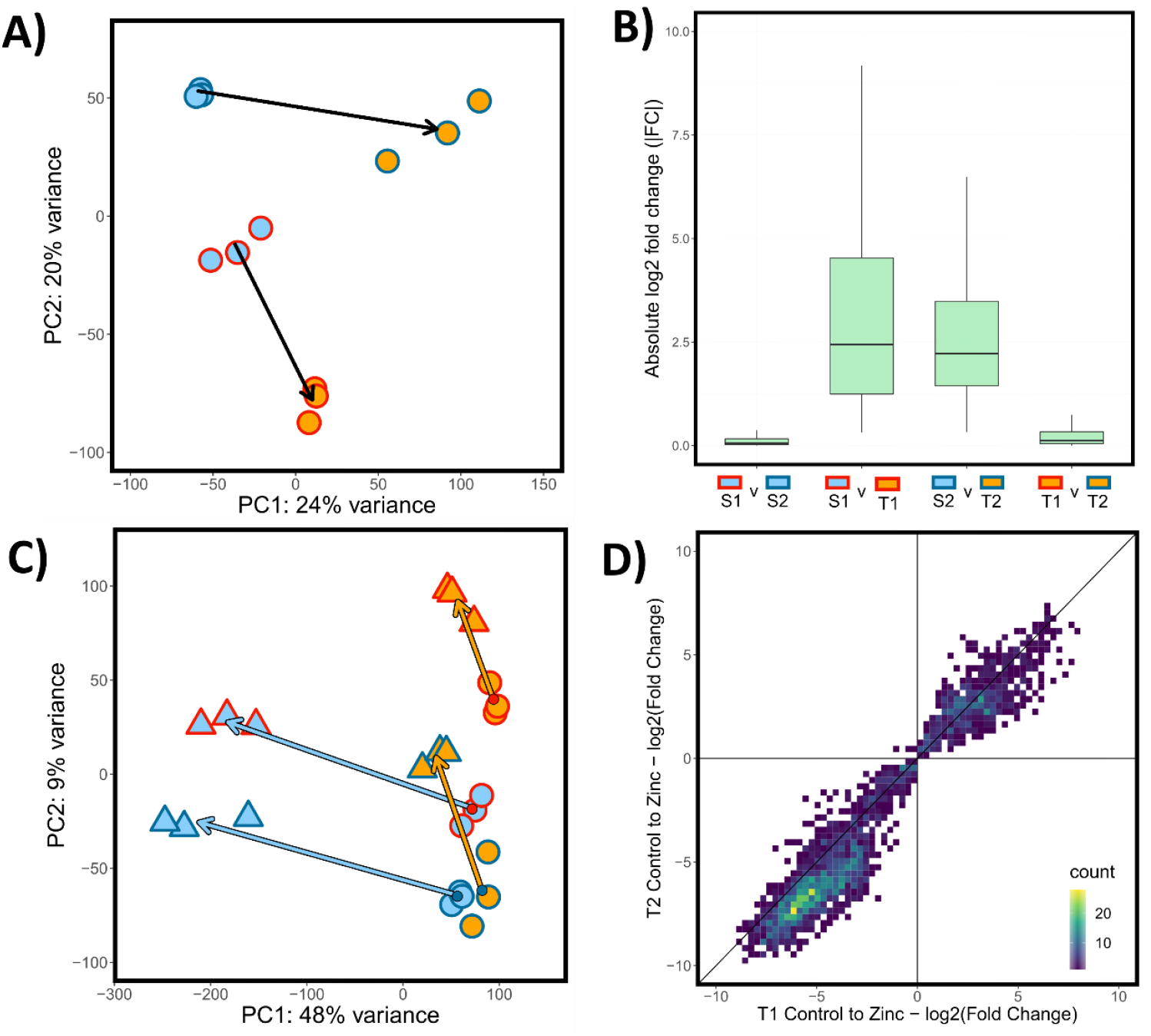
Parallel constitutive and plastic changes in tolerant populations. A) Principal components analysis (PCA) of variance-transformed counts of populations in control treatments. Point fill corresponds to population zinc tolerance (orange = tolerant, blue = sensitive), point border corresponds to geographic pair (red = 1, dark blue = 2). Arrows are drawn from the centroid of each susceptible population (S1 and S2) to the centroid of their corresponding tolerant population (T1 and T2 respectively). B) Boxplots of absolute values of log2 transformed fold changes (|FC|; y axis) between pairs of populations (x axis) in the control treatment for CEC genes (i.e., 413 genes showing parallel expression differences in the control treatment). Points above 97.25th and below 2.25th percentiles not plotted. C) PCA of variance-transformed counts of all populations in both treatments across all genes. Circles correspond to control treatment, triangles to zinc treatment. Point fill and border as in panel 2A. Arrows are drawn from the centroid of each population in the control treatment to the centroid of each population in the zinc treatment (arrowhead). Arrow fill corresponds to zinc tolerance level and arrow base point fill corresponds to geographic pair (colours as in Fig. 2A). D) Heatmap of log2 transformed shrunken fold changes between control and zinc treatments for genes that were differentially expressed between control and zinc in both T1 (x-axis) and T2 (y-axis; i.e. DP genes).

However, 2,198 and 4,448 genes were differentially expressed in control conditions between T1 and S1, and T2 and S2 respectively, of which 566 were shared (Fig. S2D). We categorised 413 of these shared genes as displaying parallel constitutive evolutionary changes of expression (CEC genes); these were differentially expressed in both tolerant-sensitive pairs *and* had expression differences in the same direction (i.e., increased or decreased expression in both T1 vs. S1 and T2 vs. S2). Genes with expression shifts in the same direction are more likely to be the result of parallel adaptation to similar selection pressures across the mines (Fig. S3). With RNA-seq data, the extent to which pairs of populations exhibit similar gene expression levels can be quantified by comparing the absolute per-gene log2 transformed shrunken fold-changes (FC) between them (see Methods for rationale). A small median |FC| for a gene set indicates high similarity between a pair of populations in the expression levels of those genes. In control conditions, transcriptome-wide expression values of tolerant populations were less similar than the coastal populations were to each other - (|FC|_S1-S2_ = 0.032 vs |FC|_T1-T2_ = 0.098; p-value < 2.2 x 10^-16^; Pairwise Wilcoxon Rank Sum Test; Fig. S4). The CEC genes had similar expression values in sensitive populations (CEC|FC|_S1-S2_ = 0.055), but expression was also highly similar in tolerant populations (CEC|FC|_T1-T2_ = 0.12), despite substantial expression divergence and genome-wide genetic differentiation from the nearest coastal populations (Fig. 1B; mean FST_T-S_ = 0.36; 38). In other words, for the 413 CEC genes, parallel evolution in mine populations produced expression similarity comparable to that observed between sensitive populations - which is the product of shared ancestry, gene flow, drift and selection.

Unlike in the control treatment, there was a high degree of parallelism in the response of tolerant populations to zinc treatment across the whole transcriptome (Fig. 1C). The evolutionary trajectories from sensitive to tolerant populations in the zinc are almost completely parallel, as are the tolerant populations responses to zinc relative to the control. This suggests that genes with significant expression responses to zinc in both tolerant populations are likely to play some role zinc tolerance. Of the 2,957 and 4,837 genes displaying expression differences between treatments in T1 and T2 respectively, 2,475 were shared (Figure S2A). These shared genes had highly correlated expression shifts (log2 fold changes between treatments; linear model slope = 1.0773, p <2.2 x-10^-16,^ adjusted R^2^ = 0.936, Fig. 1D). 2,472 of these genes consistently displayed derived plasticity (DP genes; i.e., – they were differentially expressed between treatments in *both* populations, with expression shifting in the same direction). Many of these DP genes (82%) were also differentially expressed between treatments in both susceptible populations and may constitute a stress response that is represent a partially inherited from their coastal ancestors. Nevertheless, there were also convergent changes in expression levels in these genes between tolerant populations. Expression profiles for DP genes were similar across all populations in the control treatment (as summarised using PCA; Fig. S5), but, when exposed to zinc DP gene expression in tolerant and sensitive plants diverged (Fig. S5). Unlike in the susceptible populations, similarity between tolerant populations in DP genes increased in the zinc compared to the control (Fig. S6), consistent with previous studies indicating that phenotypic plasticity can result in increased phenotypic parallelism (Oke *et al*., 2016).

There were six times as many genes with derived plasticity (DP genes) as with constitutive differences between ecotypes (CEC genes). In the literature, there is significant variability across taxa in the ratios of constitutive to plastic differences associated with local adaptation (Josephs *et al*., 2015; Heckel *et al*., 2016; Gugger *et al*., 2017; Passow *et al*., 2017; Feiner *et al*., 2018; Gould *et al*., 2018; Swaegers *et al*., 2020; Fischer *et al*., 2021). This may be a function of the degree to which a stressor varies temporally and spatially within a habitat. Soil metal content does vary considerably over small spatial scales and the sessile nature of *S. uniflora* may make plastic responses that can be tweaked to the precise local zinc concentrations more advantageous than constitutive changes to expression (Baker, 1974; Deram *et al*., 2006; Bidar *et al*., 2009). Only 26 genes were categorised as both CEC and DP – a relatively small overlap compared to studies in animals, which have mainly found that large proportions of plastic genes differ between-ecotypes even in the absence of the stressor (Gleason & Burton, 2015; Li *et al*., 2018). Partial upregulation in benign conditions may be beneficial when environmental stressors vary rapidly over time (Rivera *et al*., 2021), which could be the case for stressors in studied animals, but is unlikely to be the case for soil metal concentrations encountered by sessile plants.

Overall, these results suggest that highly parallel patterns of differential gene expression across evolutionary replicates can be acquired very early in adaptation and over very short timescales. This is true for both the identity of the genes and the magnitude of expression shifts. Previous experimental evolution studies in *Drosophila, Tribolium* and *Ipomoea* have demonstrated the evolution of gene expression plasticity in response to heterogenous environments within 22–130 generations (Huang & Agrawal, 2016; Koch & Guillaume, 2020; Mallard *et al*., 2020; Josephs *et al*., 2021). We demonstrate that this can also occur in wild plant populations in comparable timeframes and is repeatable between independent colonisations of a novel habitat.

### Ancestral plasticity is generally reversed during adaptation

To understand the relationship between ancestral plasticity and adaptation, a well-established approach is to investigate mean differences in gene expression between ancestral populations in their home/control environment (L_o_), in a new environment (L_p_), and in adapted populations in the new environment [L_a_; see Fig. 2B-E; (Ghalambor *et al*., 2015; Ho & Zhang, 2018, 2019; Fischer *et al*., 2021)]. To make inferences about the role of ancestral plasticity during adaptation, we can compare the direction and magnitude of the initial plastic response of an ancestral population when it is exposed to a new environment (ancestral plasticity/plastic change, PC = L_p_-L_o_) with the subsequent change in expression between the ancestral population, and an adapted population, in the new environment [evolutionary change, EC = L_p_-L_o_; (Ghalambor *et al*., 2015; Ho & Zhang, 2018)]. The relationship between PC and EC (i.e. the evolutionary response to ancestral plasticity) can be characterised in three ways: i) “reinforcement”, where the initial PC and subsequent EC both move expression in the same direction towards the new optimum (Fig. 2A, B); (ii) “overshooting” where PC takes expression beyond the new optimum and EC then adjusts expression in the opposite direction, (Fig. 2A, C); and iii) “reversions” where the new optimum is closer to the level of the ancestor in its home environment, so EC largely counteracts the change observed in PC (Fig. 2A, D-E). During both reinforcement and overshooting, the ancestral PC moves expression closer to the new optimum, so both can be interpreted as ancestral plasticity facilitating adaptation to the new environment. Conversely, reversions are likely to be the outcome when ancestral plasticity is maladaptive.

**Figure 2:**
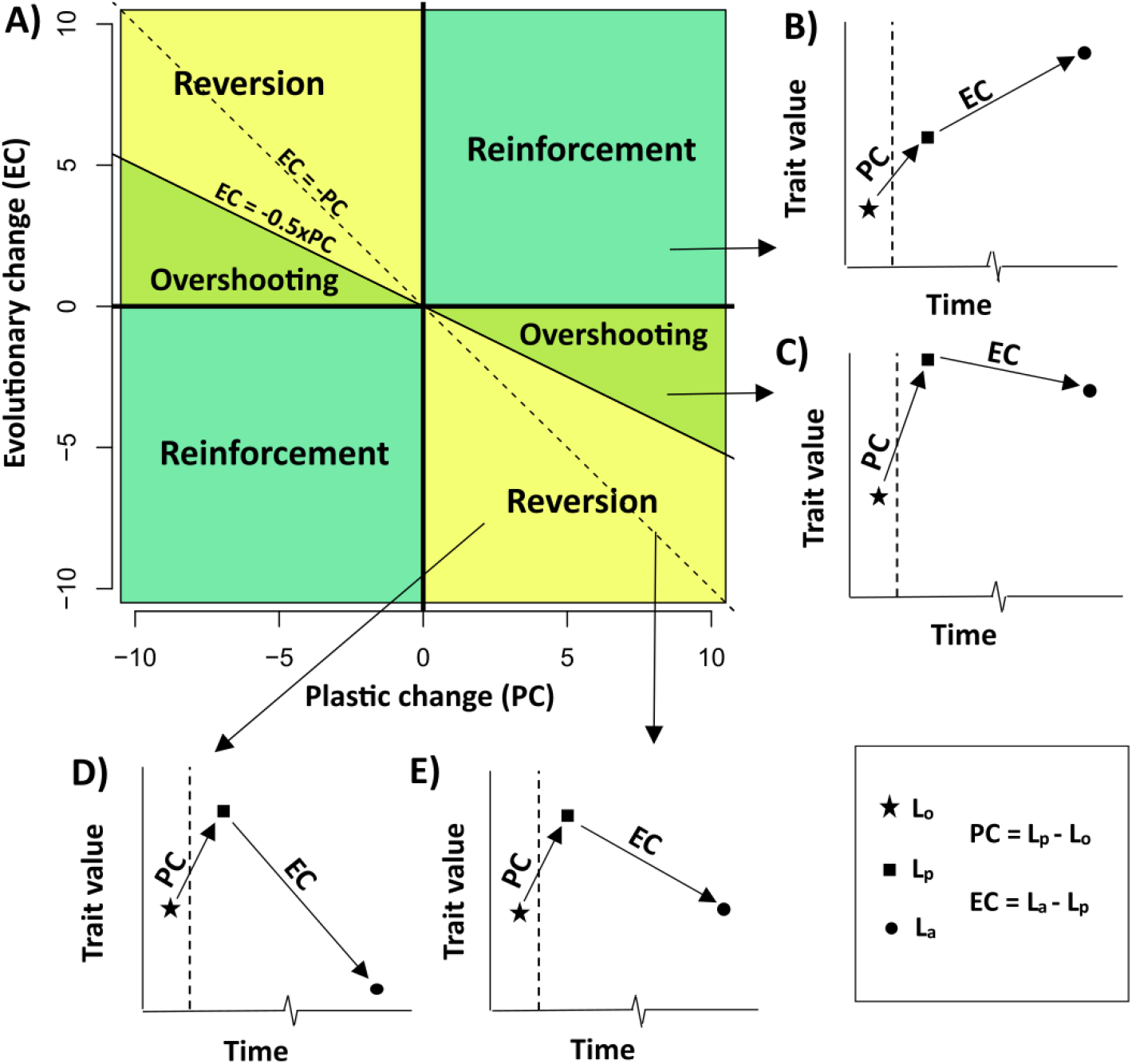
Conceptual Overview of Evolutionary Responses to Ancestral Plasticity. When an ancestral population reaches a novel environment, an immediate plastic change (PC) moves the trait from an initial value of L_o_ in the old environment to L_p_ in the new environment. As populations adapt over time, a further evolutionary change (EC) shift L_p_ to a new value of L_a_. A) The evolutionary response to ancestral plasticity can be divided into three categories depending on the values of PC and EC. B-E) Cartoon representations of scenarios – dashed line represents transition from ancestral to novel environment and associated trait shift, PC. B) Reinforcement occurs when the subsequent EC is in the same direction as PC. C) Overshooting occurs when PC has moved the trait value closer to the new optimum (i.e., L_a_ is closer to L_p_ than L_o_). In this scenario, EC is in the opposite direction to PC, but |EC| < 0.5*|PC|. D-E) Reversion occurs when the optimum in the new habitat is nearer to the value of the unstressed ancestor in its home environment then the ancestor’s response (i.e., L_a_ is closer to L_o_ than L_p_), so EC is in the opposite direction to PC, but |EC| < 0.5*|PC|. Reversion can include the restoration of the ancestral state in the old environment (|EC| = |PC|) or move beyond this value in the opposite direction (|EC|>|PC|). Reinforcement and overshooting suggest that ancestral plasticity was adaptive, whereas reversion indicates it was maladaptive.

We evaluated the degree of reversion, reinforcement and overshooting in our transcriptome dataset. To avoid spurious assignment to these categories resulting from very small expression changes, only genes showing substantial changes in PC and EC (|PC| & EC| > 0.2*L_o_; 76.7% of all genes;) were placed into these three categories (Fig. 2A), as in previous studies (Ghalambor *et al*., 2015; Ho & Zhang, 2018; Fischer *et al*., 2021). We first considered these patterns transcriptome-wide, to establish the general pattern of evolutionary responses to ancestral plasticity, regardless of these genes’ role in conferring adaptation. Across the entire transcriptome, 87.7% of genes showed reversion, with only 6.9% showing reinforcement and 5.3% overshooting. This indicates that, in the vast majority of cases, ancestral plasticity does not move expression closer to the new optimum (Figs. 3A, S7). Ho & Zhang (2019) identified that reversions may be overrepresented due to the presence of L_p_ in calculations of PC and EC (Fig. 2), recommending parametric bootstrapping to reduce this bias. Bootstrapping (see Methods) produced very similar proportions of reversions, overshooting and reinforcement across all gene sets, so this bias had minimal impact on our findings (Table S3). Our transcriptome-wide results are consistent with previous studies in animals and microorganisms which generally find that reversion is dominant (Ho & Zhang, 2018; Koch & Guillaume, 2020; Swaegers *et al*., 2020). However, the subsequent evolutionary response (EC) in many of these genes are likely to be consequences of adaptation (reduced stress/transcriptional disruption in mine populations due to their zinc tolerance, resulting in reversion of ancestral stress responses) rather than it causes (i.e., not directly involved in conferring zinc tolerance).

**Figure 3:**
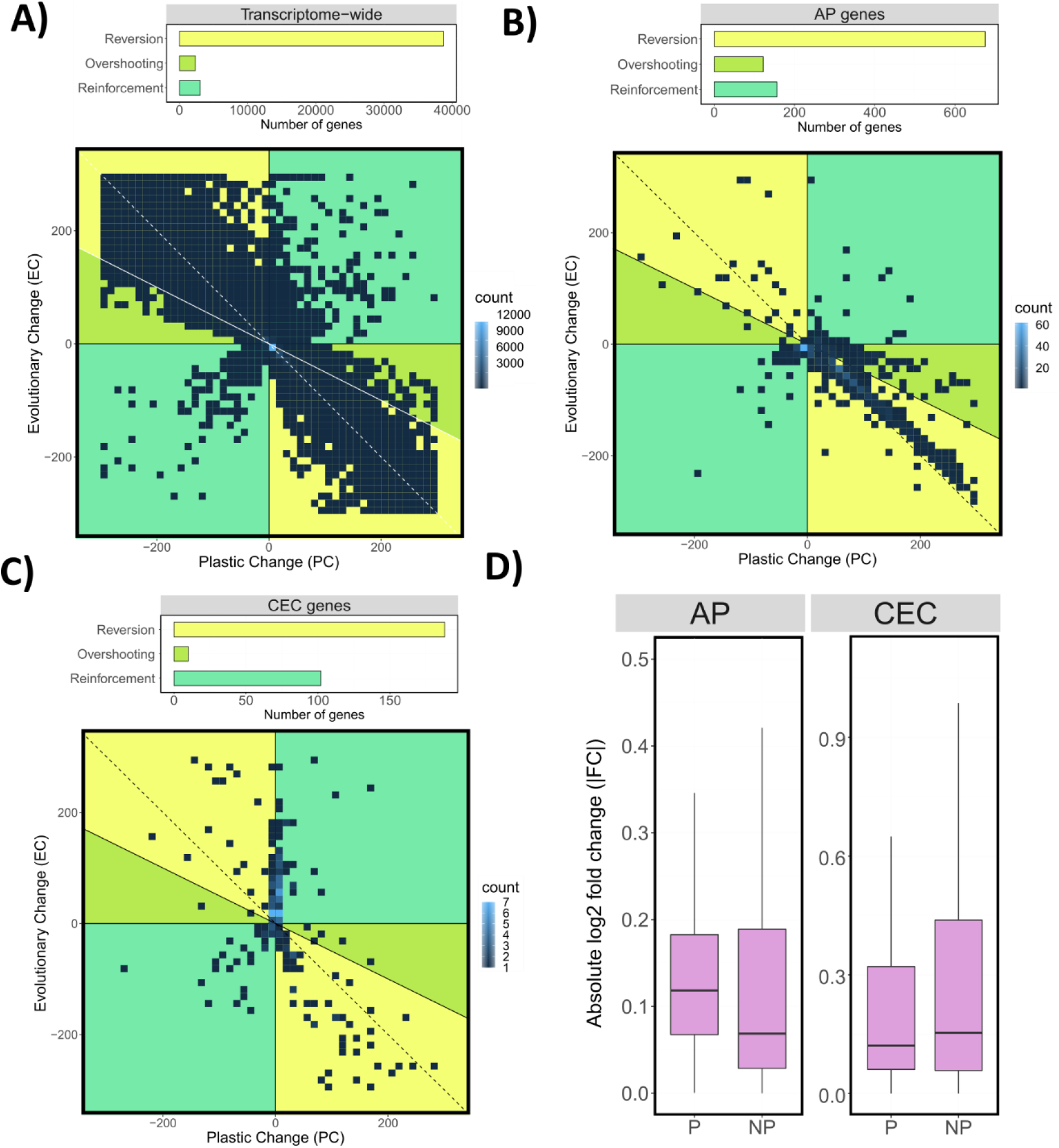
Impact of ancestral plasticity on adaptive evolution and expression convergence. For each of A) the entire transcriptome, B) derived plasticity (DP) genes, and C) genes with constitutive expression differences (CEC): i) barplots displaying numbers of genes displaying reversion, overshooting and reinforcement; and ii) heatmaps of plastic change (PC) vs. evolutionary change (EC) for each gene. Plots display at least 50% of the genes in each category (see Fig. S6 for plots of entire datasets). D) Boxplots of absolute values of log2 transformed fold changes (|FC|; y axis) between tolerant populations in the zinc for genes with derived plasticity (DP genes) or constitutive changes (CEC genes), and either substantial ancestral plasticity (P) or no ancestral plasticity (NP).

Genes that plausibly have a role in repeated adaptation to zinc contamination (DP and CEC genes) make up only 5.0% of the transcriptome. The vast majority of genes displaying substantial PC and EC across the transcriptome undergo high stress responses in sensitive plants in zinc and remain at unstressed levels in tolerant populations in the zinc treatment. Examining the evolutionary response to ancestral plasticity across the transcriptome provides an indication of the overall likelihood that an ancestral plastic response moves expression closer to the new optimum in the new zinc-contaminated environment - which our results indicate is low, as evidenced by the large number of subsequent evolutionary reversions (Fig. 3A). However, it does not identify whether this likelihood increases for genes directly involved in adaptation, which is arguably more informative in understanding the role of plasticity in adaptation to new environments.

### Ancestral plasticity less likely to be reversed in adaptive genes

To understand whether ancestral plasticity facilitates adaptive evolution, we considered the proportion of genes undergoing reversion, reinforcement and overshooting in the DP and CEC gene sets. Among DP genes showing substantial PC and EC (39% of the total), 70.1% underwent reversion (i.e., the ancestral plastic response took expression further away from the new optimum), 16.3% experienced reinforcement and 12.8% overshooting (Figs. 3B, S7). These results indicate that even among genes likely to be directly involved in zinc tolerance, most evolutionary responses involve reversion of the plastic response. However, significantly more instances of the ancestral plastic response move expression closer to the new optimum than in the transcriptome as a whole (reversion + overshooting = 29.1% for DP vs. 12.2% for the whole transcriptome, p < 2.2*10^-16^ binomial two-sided test). Few DP genes display low PC and high EC (Fig. 3B), which indicates that it is difficult/rare to evolve plasticity *de novo* during adaptation, without existing ancestral plasticity. This shows that ancestral plasticity plays an important role in colonisation of new environments with some plastic responses being enhanced or only partially reversed during adaptation. Furthermore, a majority of all DP genes (53.0%) display substantial PC but EC below the threshold value, which indicates that a significant proportion of the adaptive response involves ancestral plastic responses which are not substantially altered following adaptation (i.e., they are pre-adaptations).

Adaptation to zinc contamination has also produced constitutive gene expression differences between tolerant and sensitive populations that persist even in the absence of zinc (CEC genes). Ancestral plasticity may facilitate the evolution of differences by moving expression closer to the new optimum, which could then be canalised to produce constitutive adaptive changes (Heckel *et al*., 2016). Among CEC genes showing substantial PC and EC (73% of the total), only 62.7% show signs of reversion, with 34.0% undergoing reinforcement and 3.3% overshooting (Figs. 3C, S7). This is significantly higher than in either DP genes (34.0% vs. 16.3%, p < 9.3×10^-14; Two-Tailed Binomial Test) or transcriptome-wide (34.0% vs. 6.9%, p <2.2*10^-16; Two-Tailed Binomial Test). In most cases, the PC of CEC genes is small relative to EC (Fig 3C); high plasticity is more likely to be deleterious in large-effect genes, whereas lower levels of plasticity may afford individuals sufficiently high fitness to colonise the new environment without incurring fitness costs in other situations. The increase in reinforcement among CEC genes points to canalisation of enhanced ancestral plastic responses making an important contribution to the evolution of constitutive changes.

Why is there a difference in reinforcement between DP and CEC genes? For CEC genes, there may be little cost to maintaining expression at extreme values in the face of such strong selective forces, even when zinc levels are low. If this is the case, constitutively expressed genes may be expressed at values commensurate with the most extreme environment that could plausibly be encountered, which is then consistent with higher levels of reinforcement in CEC genes. Conversely, DP genes may carry a fitness cost for being expressed at an inappropriate/inaccurate level for a given concentration of zinc and have retained, or fine-tuned, the ancestral level of plasticity. Consistent with fine-tuning of the ancestral response DP genes, where ancestral plasticity took expression closer to the adapted level, there is an approximately equal frequency of reinforcement and overshooting.

Other studies have looked for a significant role for ancestral plasticity in producing constitutive expression differences by establishing a positive correlation between ancestral plasticity (which they define as L_p_/L_o_) and evolutionary change in control conditions [defined as L_c_/L_o_, where L_c_ is the level of the adapted population in the ancestral environment (Josephs *et al*., 2021; Bittner *et al*., 2021)]. However, the common denominator of L_o_ in both variables would tend to produce a positive correlation (Kenney, 1982), potentially making these results unreliable. Ghalambor et al. (2015) found most constitutive differences had evolutionary changes in the opposite direction to ancestral plasticity (reversion and overshooting were not distinguished), but whether there was an increase compared to the transcriptome-wide pattern was not assessed. Here, we demonstrated that although most ancestral plasticity is maladaptive, ancestral plasticity that does move expression closer to the new optimum contributes significantly to adaptation.

### Ancestral plasticity not necessary for substantial gene expression convergence

Given this evidence of ancestral plasticity contributing to adaptation, the question of its importance for parallelism in adaptation arises. As demonstrated above, ancestral plasticity that moves expression closer to the new optimum (reinforcement and overshooting) is overrepresented in gene sets displaying signs of parallel changes compared to the transcriptome as a whole. Such plasticity may also increase the propensity of genes to be repeatedly recruited during adaptation. Unlike the shared CEC genes, which had relatively low rates of reversion (62.7%), genes differentially expressed in the control in only one population pair, were more likely to show reversion (77.6% and 81.8% respectively; Table S4). Similarly, genes with between-condition changes in only T1 or T2 were also more likely to show reversion (89.9% and 77.6% respectively; Table S4) than genes displaying this behaviour in both (i.e., DP genes; 70.8%). In other words, genes repeatedly recruited during adaptation are more likely to have had ancestral plasticity that moved expression closer to the new optimum, than those that were only recruited in one event.

In addition to affecting the probability of gene recruitment during adaptation, ancestral plasticity may also affect the degree of expression convergence in repeatedly recruited genes. For DP genes with no substantial ancestral plasticity (|PC| < 0.2*L_o_), expression values in tolerant populations in zinc were marginally more similar than for DP genes with ancestral plasticity (|FC|_NOPLAST_ = 0.069, |FC|_PLAST_ = 0.12, p = 0.035, Pairwise Wilcoxon Rank Sum Test; Fig. 3D). CEC genes with no substantial ancestral plasticity developed expression similarity equal to that of CEC genes with substantial ancestral plasticity (|FC|_NOPLAST_ = 0.15, |FC|_PLAST_ = 0.12, p = 0.41, Fig. 3D). This indicates that genes lacking ancestral plasticity can rapidly evolve plastic responses with comparable levels of expression convergence to those genes where plasticity is at least partly inherited from zinc-sensitive ancestors. Overall, the results show that ancestral plasticity can facilitate the repeated recruitment of genes during adaptation, but it does not facilitate greater convergence in expression levels of these genes.

### Convergent zinc tolerance pathways

Examining sets of shared genes with expression patterns consistent with a role of adaptation sheds light on the mechanisms underlying zinc tolerance. CEC genes that were upregulated by tolerant plants relative to sensitive plants included homologs of *A. thaliana zinc transporter 1* ZIP1, which mediates the uptake of zinc from the rhizosphere (Grotz *et al*., 1998), *heavy metal atpase 2* [HMA2, a plasma membrane protein that transports zinc from cells; (Hussain *et al*., 2004; Eren & Argüello, 2004)] and *metal tolerance protein 1* [MTP1, which sequesters zinc into vacuoles and controls zinc accumulation in roots; (Van Zaal *et al*., 1999; Kobae *et al*., 2004)] These are upregulated in zinc hyperaccumulators such as *Arabidopsis halleri* (Assuncao *et al*., 2001) and when overexpressed confer increased metal accumulation and tolerance (Van Zaal *et al*., 1999; Verret *et al*., 2004; Das *et al*., 2016). The function of these genes is consistent with increased zinc accumulation in the roots of zinc-tolerant *S. uniflora* populations (Baker, 1978; Papadopulos *et al*., 2021). The DP genes that were upregulated in response to zinc were enriched for GO terms including “toxin catabolic process” and “glutathione metabolic process” (Table S5). These terms were associated with the same four genes, which are homologs to *A. thaliana* glutathione-s-transferases [GSTs; which have an important role in xenobiotic detoxification (Martinoia *et al*., 1993)]. Overexpression of GSTs results in enhanced zinc and cadmium tolerance (Liu *et al*., 2013; Zhang *et al*., 2019). This indicates that genes which have been repeatedly recruited for a role in zinc tolerance across multiple species (Singh *et al*., 2016) have also undergone repeated gene expression changes in zinc-tolerant populations over a few hundred generations.

### Conclusions

Highly parallel gene expression phenotypes have evolved in *S. uniflora* during the repeated colonisation of zinc-contaminated mines, despite the short timescales involved and a lack of gene flow between the tolerant populations (Papadopulos *et al*., 2021). We show that genes displaying beneficial patterns of ancestral plasticity are overrepresented in these highly parallel gene sets, confirming a role for ancestral plasticity in facilitating repeated adaptation to novel environments. The results of our experiment and others confirm that most ancestral plasticity is non-adaptive (Ho & Zhang, 2018; Koch & Guillaume, 2020; Fischer *et al*., 2021). Nevertheless, the considerable proportion of genetically based adaptive differences that co-opt ancestral plastic responses, suggests that it is a major force in rapid adaptation. Despite a role for ancestral plasticity in enhancing the recruitment of genes, it does not result in an increased level of phenotypic convergence at the level of gene expression compared to genes showing no significant ancestral plasticity. In other words, ancestral plasticity only facilitates parallel evolution at certain levels of biological organisation. Overall, our results indicate that genetic assimilation and modification of ancestral plastic responses play an important role in adaptation to novel environments and may be partially responsible for parallelism in gene expression during local adaptation.

## Methods

### 1. Plant materials and experimental procedure

Populations T1, S1, T2 and S2 correspond to WWA-M, WWA-C, ENG-M and ENG-C in Papadopulos et al., 2021 - seeds were collected as described in that study. Seeds from three individuals per population were germinated and cuttings propagated at ten weeks (See SI Appendix for conditions). Cuttings were transferred to deep water culture tanks containing dilute Hoagland’s solution. Cuttings from each individual were included in each tank and there was approximately equal representation of populations per tank. After one week of acclimation, the hydroponic solution was replaced with fresh solution in 50% of tanks (control treatment) and solution adjusted to 600μM ZnSO_4_ solution in the remaining 50% (zinc treatment). Eight days later, roots from each individual cutting were flash frozen in liquid nitrogen and stored at −80°C. For each individual within a treatment, roots of three cuttings were pooled, homogenised and RNA extracted using a Qiagen RNeasy Plant Mini Kit (see SI Appendix for full experimental and extraction conditions). RNA-seq libraries were sequenced at the Beijing Genomics Institute in Hong Kong on a BGISEQ500 with 100bp paired-end reads (mean insert size 161bp), producing 25.1-26.0M read pairs per sample (Table S6).

### 2. Transcriptome assembly and transcript quantification

After quality control and trimming of sequencing reads (see SI appendix for details), *de novo* transcriptome assembly was performed using Trinity v2.10.0 (Haas *et al*., 2013) using data from one individual per population per treatment (Table S6). After filtering (see SI Appendix for details) 57,541 genes were retained for downstream analysis. Completeness was assessed using the Eudicots dataset in BUSCO v.4.0.5 (Seppey *et al*., 2019) - 75% complete (72.2% single copy, 2.8% duplicated), 8.4% fragmented, 16.6% missing. Transcripts were annotated using hmmer-3.3 (Mistry *et al*., 2013) blastp and trinotate v3.2.1 (Bryant *et al*., 2017; see SI Appendix for details).

### 3. Differential gene expression

Abundance estimates for transcripts were summarised at the gene level using tximport v.1.4.2 (Soneson *et al*., 2015). Gene expression analysis was performed using DESeq2 v1.26.0 (Love *et al*., 2014). Genes with low counts (<10) across all samples were removed. Variance-stabilising transformed counts for 57,476 genes across all conditions were calculated and used in downstream analysis. Principal components analysis of these counts for i) all genes in control conditions (Figure 1A), ii) all genes across all conditions (Figure 1D) and iii) for DP genes (Figure S4) were calculated using the R prcomp function.

Genes differentially expressed between two populations within a treatment (control or zinc) were identified using DESeq2’s in-built models with a single combined factor for population + condition (adjusted p-value = 0.05). Differentially expressed genes between T1 and S1, and T2 and S2 were identified in i) control and ii) zinc treatments. CEC genes were defined as those differentially expressed between both T1 and S1 in the control, and T2 and S2 in the control, in the same direction (i.e. both increasing, or decreasing, in T1 relative to S1 and T2 relative to S2). For between-treatment, within-population comparisons, a model with terms “~ Population + Population:Individual + Population:Condition” was fitted to account for individual-specific variation which could be accounted for due to the use of cloned individuals. Genes differentially expressed between control and zinc were identified for S1, S2, T1 and T2. DP genes were defined as those differentially expressed between conditions in both T1 and T2 in the same direction (i.e. both increasing, or both decreasing, from control to zinc treatment). The significance of overlaps between sets of differentially expressed genes was determined using Fisher’s Exact test (Supplementary Table S7). Gene Ontology enrichment analysis of gene sets was performed using GOseq v1.38.0 (Young *et al*., 2010) with a false discovery rate of 0.05.

Quantification of fold changes of genes between populations and/or treatments used empirical bayes shrinkage, calculated with the lfcShrink() function in DESeq2 (Stephens, 2017). Values of |FC| were calculated for each gene as the absolute log2 fold change between pairs of population/treatment groups (e.g. T1 and T2 in the zinc) for a given set of genes. The sign of the log2 fold change depends on the order of comparisons being made (e.g. a value of +1 between T1 and T2 is equivalent of −1 between T2 and T1); the absolute value must be taken to meaningfully summarise the difference in expression levels (e.g. the mean of −2 and +2 would be lower than that of 0.5 and 0.6). The median was used to summarise the values of |FC| as their distribution is highly skewed. Pairwise Wilcoxon signed-rank tests with Benjamini-Hochberg correction were used to detect significant differences in the distributions of |FC| between different pairs of population/treatment groups.

### 4. Classifying responses to ancestral plasticity

To classify evolutionary responses to ancestral plasticity in the transcriptome-wide, DP and CEC gene sets, the following parameters were calculated for each gene: L_o_ – mean expression value across S1 and S2 in control; L_p_ – mean expression value across S1 and S2 in zinc; La – mean expression value across T1 and T2 in the zinc. These were used to calculate the initial plastic change (PC = L_a_-L_p_) and subsequent evolutionary change (EC = L_p_ – L_o_) as in Ghalambor *et al*. (2015) for each gene. Only genes having substantial plastic and evolutionary change (defined as |EC| and |PC| > 0.2*L_o_), were assigned as undergoing reversion, reinforcement or plasticity – very small values of EC or PC due to measurement error would lead to spurious assignment of genes to categories (Ho & Zhang, 2018). Genes were assigned to one of three categories of evolutionary response to ancestral plasticity: i) Reinforcement: if EC*PC > 0; ii) Overshooting: if EC*PC < 0 and |EC| < 0.5*|PC|; or iii) Reversion: if EC*PC < 0 and |EC| > 0.5*|PC| (Koch & Guillaume, 2020). Significant differences in the relative proportions of these categories between sets of genes (e.g. CEC genes compared to the transcriptome as a whole) were assessed using a two-tailed binomial test. Parametric bootstrapping of gene assignment to these categories following recommendations in Ho & Zhang (2019) was implemented in R and repeated 100 times per gene (see SI Appendix for details) but showed little difference from non-bootstrapped results (Table S3), values for the latter were used throughout. For genes showing DP/CEC expression patterns but in T1/S1 or T2/S2 only, values of L_o_, L_p_, L_a_, EC and PC were only calculated using the samples from T1/S1 and T2/S2 separately (Table S4) and categorized based on these values. Assignment of categories for transcriptome-wide, CEC and DP genes were also calculated using T1/S1 and T2/S2 separately; these did not differ substantially between evolutionary replicates or the combined calculations (Table S4).

### 5. Genotyping

For genotyping, cleaned reads were mapped to the transcriptome using HISAT2 v2.2.1 (Kim *et al*., 2019). Genotypes were called using bcftools (See SI Appendix for details). A phylogenetic tree was constructed based on 24,982 SNPs using SNPhylo v20140701 (Lee *et al*., 2014).

## Materials and Data

Raw reads are uploaded to NCBI SRA, accession number PRJNA706929. The *de-novo* assembled transcriptome is uploaded to NCBI Genbank TSA, accession number GFXXXXX.

## Author Contributions

ASTP conceived and supervised the research. ASTP and DW designed the experiment with contributions from all authors. ASTP, RJS, JL conducted fieldwork. DW and JAH conducted experimental and laboratory work. DW analysed the data with contributions from ASTP, OGO, ARE, LTD and AJH. DW and ASTP wrote the manuscript and all authors commented on the final version.

## Acknowledgements

We thank Natural Environment Research Council (NERC) for funding (NE/R001081/1), Aaron Comeault and Michael Chester for valuable discussion, Nicholas Welsby and Wendy Grail for laboratory support, and Llinos Hughes and Mark Hughes for greenhouse support. LTD is supported by a Natural Environment Research Council Independent Research Fellowship (NE/T011025/1).

## References

Assuncao AGL, Martins PDC, De Folter S, Vooijs R, Schat H, Aarts MGM. 2001. Elevated expression of metal transporter genes in three accessions of the metal hyperaccumulator Thlaspi caerulescens. Plant, Cell and Environment 24: 217–226.

Baker AJM. 1974. Heavy metal tolerance and population differentiation in Silene maritima With.

Baker AJM. 1978. Ecophysiological Aspects of Zinc Tolerance in Silene maritima With. New Phytologist 80: 635–642.

Baldwin JM. 1896. A New Factor in Evolution. The American Naturalist 30: 441–451.

Bidar G, Pruvot C, Garçon G, Verdin A, Shirali P, Douay F. 2009. Seasonal and annual variations of metal uptake, bioaccumulation, and toxicity in Trifolium repens and Lolium perenne growing in a heavy metal-contaminated field. Environmental Science and Pollution Research 16: 42–53.

Bittner NKJ, Mack KL, Nachman MW. 2021. Gene expression plasticity and desert adaptation in house mice*. Evolution 75: 1477–1491.

Bolnick DI, Barrett RDH, Oke KB, Rennison DJ, Stuart YE. 2018. (Non)Parallel Evolution. Annual Review of Ecology, Evolution, and Systematics 49: 303–330.

Bryant DM, Johnson K, DiTommaso T, Tickle T, Couger MB, Payzin-Dogru D, Lee TJ, Leigh ND, Kuo TH, Davis FG, et al. 2017. A Tissue-Mapped Axolotl De Novo Transcriptome Enables Identification of Limb Regeneration Factors. Cell Reports 18: 762–776.

Christin PA, Weinreich DM, Besnard G. 2010. Causes and evolutionary significance of genetic convergence. Trends in Genetics 26: 400–405.

Conway Morris S. 2003. Life’s Solution: Inevitable Humans in a Lonely Universe - Simon Conway Morris - Google Books. Cambridge: Cambridge University Press.

Das N, Bhattacharya S, Maiti MK. 2016. Enhanced cadmium accumulation and tolerance in transgenic tobacco overexpressing rice metal tolerance protein gene OsMTP1 is promising for phytoremediation. Plant Physiology and Biochemistry 105: 297–309.

Deram A, Denayer FO, Petit D, Van Haluwyn C. 2006. Seasonal variations of cadmium and zinc in Arrhenatherum elatius, a perennial grass species from highly contaminated soils. Environmental Pollution 140: 62–70.

Draghi JA, Whitlock MC. 2012. Phenotypic plasticity facilitates mutational variance, genetic variance, and evolvability along the major axis of environmental variation. Evolution 66: 2891–2902.

Eren E, Argüello JM. 2004. Arabidopsis HMA2, a Divalent Heavy Metal-Transporting P IB-Type ATPase, Is Involved in Cytoplasmic Zn ^2+^ Homeostasis. Plant Physiology 136: 3712–3723.

Feiner N, Rago A, While GM, Uller T. 2018. Signatures of selection in embryonic transcriptomes of lizards adapting in parallel to cool climate. Evolution 72: 67–81.

Fischer EK, Song Y, Hughes KA, Zhou W, Hoke KL. 2021. Nonparallel transcriptional divergence during parallel adaptation. Molecular Ecology 30: 1516–1530.

Ghalambor CK, Hoke KL, Ruell EW, Fischer EK, Reznick DN, Hughes KA. 2015. Non-adaptive plasticity potentiates rapid adaptive evolution of gene expression in nature. Nature 525: 372–375.

Ghalambor CK, McKay JK, Carroll SP, Reznick DN. 2007. Adaptive versus non-adaptive phenotypic plasticity and the potential for contemporary adaptation in new environments. Functional Ecology 21: 394–407.

Gleason LU, Burton RS. 2015. RNA-seq reveals regional differences in transcriptome response to heat stress in the marine snail Chlorostoma funebralis. Molecular Ecology 24: 610–627.

Gould SJ. 1989. Wonderful life: The Burgess shale and the nature of history. New York City: W. W. Norton & Co.

Gould BA, Chen Y, Lowry DB. 2018. Gene regulatory divergence between locally adapted ecotypes in their native habitats. Molecular Ecology 27: 4174–4188.

Grotz N, Fox T, Connolly E, Park W, Guerinot M Lou, Eide D. 1998. Identification of a family of zinc transporter genes from Arabidopsis that respond to zinc deficiency. Proceedings of the National Academy of Sciences of the United States of America 95: 7220–7224.

Gugger PF, Peñaloza-Ramírez JM, Wright JW, Sork VL. 2017. Whole-transcriptome response to water stress in a California endemic oak, Quercus lobata. Tree Physiology 37: 632–644.

Haas BJ, Papanicolaou A, Yassour M, Grabherr M, Blood PD, Bowden J, Couger MB, Eccles D, Li B, Lieber M, et al. 2013. De novo transcript sequence reconstruction from RNA-seq using the Trinity platform for reference generation and analysis. Nature Protocols 8: 1494–1512.

Hanson D, Hu J, Hendry AP, Barrett RDH. 2017. Heritable gene expression differences between lake and stream stickleback include both parallel and antiparallel components. Heredity 119: 339–348.

Hargreaves AD, Swain MT, Hegarty MJ, Logan DW, Mulley JF. 2014. Restriction and Recruitment—Gene Duplication and the Origin and Evolution of Snake Venom Toxins. Genome Biology and Evolution 6: 2088–2095.

Heckel K von, Stephan W, Hutter S. 2016. Canalization of gene expression is a major signature of regulatory cold adaptation in temperate Drosophila melanogaster. BMC Genomics 17.

Ho WC, Zhang J. 2018. Evolutionary adaptations to new environments generally reverse plastic phenotypic changes. Nature Communications 9: 1–11.

Ho WC, Zhang J. 2019. Genetic Gene Expression Changes during Environmental Adaptations Tend to Reverse Plastic Changes Even after the Correction for Statistical Nonindependence. Molecular Biology and Evolution 36: 604–612.

Huang Y, Agrawal AF. 2016. Experimental Evolution of Gene Expression and Plasticity in Alternative Selective Regimes (DJ Begun, Ed.). PLOS Genetics 12: e1006336.

Hussain D, Haydon MJ, Wang Y, Wong E, Sherson SM, Young J, Camakaris J, Harper JF, Cobbett CS. 2004. P-type ATPase heavy metal transporters with roles in essential zinc homeostasis in arabidopsis. Plant Cell 16: 1327–1339.

Jacobs A, Carruthers M, Yurchenko A, Gordeeva N V., Alekseyev SS, Hooker O, Leong JS, Minkley DR, Rondeau EB, Koop BF, et al. 2020. Parallelism in eco-morphology and gene expression despite variable evolutionary and genomic backgrounds in a Holarctic fish. PLoS Genetics 16: e1008658.

Josephs EB, Etten ML Van, Harkess A, Platts A, Baucom RS. 2021. Adaptive and maladaptive expression plasticity underlying herbicide resistance in an agricultural weed. Evolution Letters.

Josephs EB, Lee YW, Stinchcombe JR, Wright SI. 2015. Association mapping reveals the role of purifying selection in the maintenance of genomic variation in gene expression. Proceedings of the National Academy of Sciences of the United States of America 112: 15390–15395.

Kelly M. 2019. Adaptation to climate change through genetic accommodation and assimilation of plastic phenotypes. Philosophical Transactions of the Royal Society B: Biological Sciences 374.

Kenkel CD, Matz M V. 2017. Gene expression plasticity as a mechanism of coral adaptation to a variable environment. Nature Ecology & Evolution 1: 14.

Kenney BC. 1982. Beware of spurious self-correlations!; Water Resources Research 18: 1041–1048.

Kim D, Paggi JM, Park C, Bennett C, Salzberg SL. 2019. Graph-based genome alignment and genotyping with HISAT2 and HISAT-genotype. Nature Biotechnology 37: 907–915.

Kobae Y, Uemura T, Sato MH, Ohnishi M, Mimura T, Nakagawa T, Maeshima M. 2004. Zinc Transporter of Arabidopsis thaliana AtMTP1 is Localized to Vacuolar Membranes and Implicated in Zinc Homeostasis. Plant and Cell Physiology 45: 1749–1758.

Koch EL, Guillaume F. 2020. Restoring ancestral phenotypes is a general pattern in gene expression evolution during adaptation to new environments in Tribolium castaneum. Molecular Ecology 29: 3938–3953.

Lee TH, Guo H, Wang X, Kim C, Paterson AH. 2014. SNPhylo: A pipeline to construct a phylogenetic tree from huge SNP data. BMC Genomics 15: 162.

Levis NA, Isdaner AJ, Pfennig DW. 2018. Morphological novelty emerges from pre-existing phenotypic plasticity. Nature Ecology and Evolution 2: 1289–1297.

Li L, Li A, Song K, Meng J, Guo X, Li S, Li C, De Wit P, Que H, Wu F, et al. 2018. Divergence and plasticity shape adaptive potential of the Pacific oyster. Nature Ecology and Evolution 2: 1751–1760.

Liu D, Liu Y, Rao J, Wang G, Li H, Ge F, Chen C. 2013. Overexpression of the glutathione S-transferase gene from Pyrus pyrifolia fruit improves tolerance to abiotic stress in transgenic tobacco plants. Molecular Biology 47: 515–523.

Losos JB. 2011. Convergence, adaptation, and constraint. Evolution 65: 1827–1840.

Love MI, Huber W, Anders S. 2014. Moderated estimation of fold change and dispersion for RNA-seq data with DESeq2. Genome Biology 15: 550.

Mäkinen H, Papakostas S, Vøllestad LA, Leder EH, Primmer CR. 2016. Plastic and evolutionary gene expression responses are correlated in European grayling (Thymallus thymallus) subpopulations adapted to different thermal environments. Journal of Heredity 107: 82–89.

Mallard F, Nolte V, Schlötterer C. 2020. The Evolution of Phenotypic Plasticity in Response to Temperature Stress. Genome Biology and Evolution 12: 2429–2440.

Martinoia E, Grill E, Tommasini R, Kreuz K, Amrhein N. 1993. ATP-dependent glutathione S-conjugate ‘export’ pump in the vacuolar membrane of plants. Nature 364: 247–249.

Mistry J, Finn RD, Eddy SR, Bateman A, Punta M. 2013. Challenges in homology search: HMMER3 and convergent evolution of coiled-coil regions. Nucleic Acids Research 41: e121–e121.

Moczek AP, Sultan S, Foster S, Ledón-Rettig C, Dworkin I, Nijhout HF, Abouheif E, Pfennig DW. 2011. The role of developmental plasticity in evolutionary innovation. Proceedings of the Royal Society B: Biological Sciences 278: 2705–2713.

Moreno-Villena JJ, Dunning LT, Osborne CP, Christin PA. 2018. Highly Expressed Genes Are Preferentially Co-Opted for C 4 Photosynthesis. Molecular Biology and Evolution 35: 94–106.

Oke KB, Bukhari M, Kaeuffer R, Rolshausen G, Räsänen K, Bolnick DI, Peichel CL, Hendry AP. 2016. Does plasticity enhance or dampen phenotypic parallelism? A test with three lake-stream stickleback pairs. Journal of Evolutionary Biology 29: 126–143.

Orgogozo V. 2015. Replaying the tape of life in the twenty-first century. Interface Focus 5.

Papadopulos AST, Helmstetter AJ, Osborne OG, Comeault AA, Wood DP, Straw EA, Mason L, Fay MF, Parker J, Dunning LT, et al. 2021. Rapid Parallel Adaptation to Anthropogenic Heavy Metal Pollution. Molecular Biology and Evolution 38: 3724–3736.

Parker DJ, Bast J, Jalvingh K, Dumas Z, Robinson-Rechavi M, Schwander T. 2019. Repeated evolution of asexuality involves convergent gene expression changes. Molecular Biology and Evolution 36: 350–364.

Passow CN, Henpita C, Shaw JH, Quackenbush CR, Warren WC, Schartl M, Arias-Rodriguez L, Kelley JL, Tobler M. 2017. The roles of plasticity and evolutionary change in shaping gene expression variation in natural populations of extremophile fish. Molecular Ecology 26: 6384–6399.

Rivera HE, Aichelman HE, Fifer JE, Kriefall NG, Wuitchik DM, Wuitchik SJS, Davies SW. 2021. A framework for understanding gene expression plasticity and its influence on stress tolerance. Molecular Ecology 30: 1381–1397.

Schaum E, Rost B, Millar AJ, Collins S. 2013. Variation in plastic responses of a globally distributed picoplankton species to ocean acidification. Nature Climate Change 3: 298–302.

Scoville AG, Pfrender ME. 2010. Phenotypic plasticity facilitates recurrent rapid adaptation to introduced predators. Proceedings of the National Academy of Sciences 107: 4260–4263.

Seppey M, Manni M, Zdobnov EM. 2019. BUSCO: Assessing genome assembly and annotation completeness. In: Methods in Molecular Biology. Humana Press Inc., 227–245.

Singh S, Parihar P, Singh R, Singh VP, Prasad SM. 2016. Heavy metal tolerance in plants: Role of transcriptomics, proteomics, metabolomics, and ionomics. Frontiers in Plant Science 6: 1143.

Soneson C, Love MI, Robinson MD. 2015. Differential analyses for RNA-seq: transcript-level estimates improve gene-level inferences. F1000Research 4: 1521.

Stephens M. 2017. False discovery rates: A new deal. Biostatistics 18: 275–294.

Stern DL. 2013. The genetic causes of convergent evolution.

Stern DB, Crandall KA. 2018. The Evolution of Gene Expression Underlying Vision Loss in Cave Animals. Molecular Biology and Evolution 35: 2005–2014.

Swaegers J, Spanier KI, Stoks R. 2020. Genetic compensation rather than genetic assimilation drives the evolution of plasticity in response to mild warming across latitudes in a damselfly. Molecular Ecology 29: 4823–4834.

Velotta JP, Ivy CM, Wolf CJ, Scott GR, Cheviron ZA. 2018. Maladaptive phenotypic plasticity in cardiac muscle growth is suppressed in high-altitude deer mice. Evolution 72: 2712–2727.

Verret F, Gravot A, Auroy P, Leonhardt N, David P, Nussaume L, Vavasseur A, Richaud P. 2004. Overexpression of AtHMA4 enhances root-to-shoot translocation of zinc and cadmium and plant metal tolerance. FEBS Letters 576: 306–312.

Waldvogel A, Feldmeyer B, Rolshausen G, Exposito-Alonso M, Rellstab C, Kofler R, Mock T, Schmid K, Schmitt I, Bataillon T, et al. 2020. Evolutionary genomics can improve prediction of species’ responses to climate change. Evolution Letters 4: 4–18.

Wang SP, Althoff DM. 2019. Phenotypic plasticity facilitates initial colonization of a novel environment. Evolution 73: 303–316.

Young MD, Wakefield MJ, Smyth GK, Oshlack A. 2010. Gene ontology analysis for RNA-seq: accounting for selection bias. Genome Biology 11: R14.

Van Zaal BJD, Neuteboom LW, Pinas JE, Chardonnens AN, Schat H, Verkleij JAC, Hooykaas PJJ. 1999. Overexpression of a novel Arabidopsis gene related to putative zinc-transporter genes from animals can lead to enhanced zinc resistance and accumulation. Plant Physiology 119: 1047–1055.

Zhang H, Yang J, Li W, Chen Y, Lu H, Zhao S, Li D, Wei M, Li C. 2019. PuHSFA4a enhances tolerance to excess zinc by regulating reactive oxygen species production and root development in populus. Plant Physiology 180: 2254–2271.

