## Supplementary Appendix for "Genetic assimilation of ancestral plasticity during parallel adaptation"

Daniel P. Wood[1]

Jon A. Holmberg[1]

Owen G. Osborne[1]

Andrew J. Helmstetter[2]

Luke T. Dunning[3]

Amy R. Ellison[1]

Rhian J. Smith[4]

Jackie Lighten[5]

Alexander S.T. Papadopulos[1]*

1. Molecular Ecology and Evolution Bangor, Bangor University, Environment Centre for Wales, Deiniol Road, Bangor, LL57 2UW, United Kingdom.

2.FRB-CESAB, Institut Bouisson Bertrand, Rue de l'École de Médecine, 34000 Montpellier

3. Department of Animal and Plant Sciences, University of Sheffield, Alfred Denny Building, Western Bank, Sheffield, S10 2TN, United Kingdom.

4. Royal Botanic Gardens, Kew, Richmond, Surrey, TW9 3AE.

5. College of Life and Environmental Sciences, University of Exeter, Prince of Wales Road, Exeter, EX4PS, United Kingdom.

***Corresponding author Alexander S.T. Papadopulos**

**This PDF file includes:**

Supplementary text
Figures S1 to S6
Tables S3, S6
Legends for Tables S1-S2, S4-S5
SI References

**Other supplementary materials for this manuscript include the following:**

Table S1-S2, S4-S5

**Supplementary Methods**

1. Plant materials and experimental procedure

Seeds from each of the four populations were collected as described in Papadopulos *et al.* (1). Populations T1, S1, T2 and S2 correspond to populations WWA-M, WWA-C, ENG-M and ENG-C in Papadopulos et al. (1). In October 2019 wild-collected seeds from at least three individuals per population were germinated in Erin Traditional Multipurpose Compost in 80ml cells and grown in a greenhouse (16/8 hour day/night cycle; temperature controlled 18/12 ºC, supplementary lighting automatically switched on if light levels fell below 120 μmol $m^{-2}$ $s^{-1}$ during the day) before being transplanted to 1.5L pots filled with compost and 14.7g MiracleGro Slow Release Fertiliser after five weeks**.** When the plants were ten weeks old at least six 6-8cm cuttings per plant were taken per plant and suspended in a foam discs (1 disc per individual) in a 40 Site X-Stream Aeroponic Propagator filled with 12L Ultrapure deioinised water (Barnstead D4642-33 e-pure ultra water purification system). After 2.5 weeks the propagator was refreshed with 1.92g Hoaglands No. 2 Basalt Salt Solution (Merck) dissolved in 12L ultrapure water once a week for a further three weeks to encourage root growth.

The rooted cuttings were then transferred to six hydroponics tanks. For each tank, 1.28g Hoaglands No. 2 Basal Salt Solution was dissolved in 0.8L deionised water (Fluid Science) and adjusted to pH 5.5 using 0.1M Potassium Hydroxide. This was added to an 18L opaque tank (48 x 39 x 20cm) and topped up to 8L using deionised water. Tanks were aerated using an airstone and pump. Individual cuttings in foam discs were added to 5x4 polystyrene floats, with 18-19 cuttings per tank, with cuttings from each individual present at least once in each tank and approximately equal representation of each population in each tank.

After one week of acclimation, the hydroponic solution was replaced. For three of the six tanks, fresh hydroponics solution prepared as previously described was added (control treatment). For the remaining three tanks, 1.28g of Hoaglands No2 Basal Salt solution and 2.4mL 2.0M $ZnSO_4$ solution (Sigma) were dissolved in 0.798 ml deioinised water, and adjusted to pH 5.5 as above. This was then topped up to 8L with deioinised water, giving a final $ZnSO_4$ concentration of 600µM (zinc treatment). Eight days after media replacement, the roots from each individual cutting were flash frozen in liquid nitrogen and stored on dry ice before being transferred to a -80˚C freezer. From each individual within a treatment, one cutting's roots from each of the three tanks (which were of approximately equal size) were pooled. This resulted in each of the three individuals per population being represented by a pooled sample of three cuttings (one from each of the three replicate tanks), for both the control and the zinc treatment. This approach reduced technical variation from between-cutting and between-tank differences, and allowed between-individual variation to be accounted for when comparing within populations between the two treatments.

2. RNA extraction and sequencing

Roots from each sample were homogenised on dry ice using a pestle to evenly distribute material from each of the three replicate cuttings. From this, a volume of approximately 150µL of material was added to 2.0mL Safe-Lock Tube (Eppendorf) along with a 5mm stainless steel bead (Qiagen). Samples were homogenised using a TissueLyser LT (Qiagen), the adapter of which had been chilled in a -80˚C freezer for at least 2 hours, for 1 minute at 50Hz. RNA was extracted using a Qiagen RNeasy Plant Mini Kit (Qiagen) using the standard protocol, including the optional DNase digestion step, with two 50µL elutions in RNase free water. RNA quantity and quality were checked using electrophoresis on a 1% agarose gel, a Nanodrop v2 and Qubit BR RNA Assay Kit (Thermo Fisher; Supplementary File 2 for RNA quantity and quality) using a Qubit v3. Samples were shipped on dry ice to the Beijing Genomics Institute in Hong Kong and RNA integrity measured using an Aligent Technologies 2100 Bioanalyser, with RIN ranging from 8.2-10.0 (Table S5). RNA-seq libraries were prepared according to the BGISEQ-500 RNA-Seq Library Preparation Protocol (Document NO: SOP-SS-027) with mRNA enrichment using oligo(dT)-attached magnetic beads. Libraries were sequenced on a BGISEQ500 with 100bp paired-end reads with an average insert size of 161bp. This produced 25.1-26.0M read pairs per sample (Table S5).

3. Data pre-processing, transcriptome assembly and transcript quantification.

Read quality was inspected using FastQC v0.11.8 (Andrews 2015) .The IlluQC.pl script from NGSQCToolkit v2.3.3 (3) was used to keep reads with at least 70% of the sequence having a PHRED score >20. Trimmomatic v0.39 (4) was used with settings LEADING:10 TRAILING:10 SLIDINGWINDOW:4:15 MINLEN:70 to remove leading and trailing low-quality bases with a length cut-off of 70bp. Trimmed reads were inspected for quality and adapter contamination using FastQC.

*De novo* transcriptome assembly was performed using Trinity v2.10.0 (5) using -min_kmer_cov=2 and default settings for all other parameters. The reads from control and zinc from the individual in each population with the highest total number of reads were used to construct the transcriptome (Table S5). This resulted in a transcriptome of

502,579 transcripts from 270,649 trinity "genes" (henceforth referred to as genes). Transdecoder v.5.5.0 (5) was used to predict open reading frames (ORFs) and transcripts lacking an ORF at least 100 amino acids long, which are likely to mostly be assembly artefacts, were discarded. 237,424 transcripts from 121,009 genes were retained. Reads were mapped to these transcripts using the align_and_estimate_abundance.pl utility in Trinity, using kallisto v0.46.1 (6).

To identify and remove transcripts potentially representing microorganism contaminants from the hydroponic media, for each gene the longest transcript was mapped to the *Silene latifolia* genome v1.0 [NCBI Assembly GCA_3260165.1; (7)] using BLAT v35 (8), with the database translated into six frames to protein and query three frames into protein using the arguments -q=RNAX -t=DNAX. Genes with a match >50bp were retained. To avoid discarding genes with homologs in unassembled regions of the *S. latifolia* genome (9), the translated protein sequences of the remaining transcripts were searched against the SwissProt/UniProt database [version 290; (10) using BLASTP v2.10.0 (11)]**.** Genes with a top hit to an Embryophyte with e-value $<10e^{-3}$ and match length >50bp long were also retained. This resulted in the retention of 57,541 genes for use in downstream analysis. Transcriptome completeness was assessed using BUSCO v.4.0.5 using the Eudicot dataset (12) - 75% of universal single-copy genes were assembled completely (72.2% single copy, 2.8% duplicated) with a further 8.4% fragmented, with 16.6% missing.

To annotate the transcriptome, ORFs >100 amino acids were searched against the Pfam-A v33.0 database (13) using hmmer-3.3 (14) to identify protein domains. These results along with the blastp results described above were loaded into an SQLite database using trinotate v3.2.1 (15).

**4. Parametric bootstrapping for classifying responses to ancestral plasticity**

. Parametric bootstrapping was implemented in R following the method of Ho & Zhang (23); for each gene normal distributions were generated with means $L_o$, $L_p$ and $L_a$ and sd the standard error of $L_o$, $L_p$ and $L_a$ as estimated from the data (i.e. from across the 6 individuals used to generate each parameter). Values were randomly drawn from each distribution and from these the gene was classified as reversal, overshooting or reinforcement. This was repeated 100 times per gene and genes which did not show one classification in 95 or more repeats were excluded from the analysis. The results of this analysis for the entire transcriptome, CEC and DP genes are shown in Table S3.

**4. Genotyping**

For genotyping, cleaned reads from each individual were mapped to the longest isoforms of each Trinity gene using HISAT2 v2.2.1 (24), with reads with MAPQ <20 removed using samtools v1.7 (25). Genotypes were called using bcftools with filter QUAL>20. Indels were removed, and at least 2 genotyped individuals per population were required for a site to be included, with a minimum read depth of 6 for a sample to be genotyped. Only sites in the 57,541 genes used in the gene expression analysis were included. A phylogenetic tree was constructed using SNPhylo v20140701 (26) using standard settings, which used 24,982 SNPs.

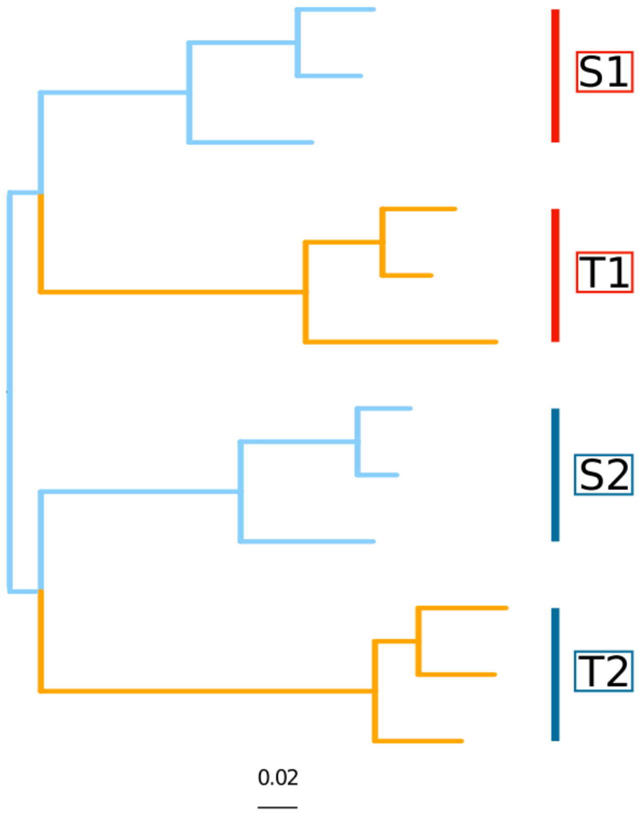


**Figure S1: Phylogenetic relationships of *S. uniflora* individuals.**

Phylogenetic relationships of the included *Silene uniflora* individuals based on 24,982 SNPs derived from the RNAseq data generated here. Mine populations are coloured orange, coastal populations coloured blue. All nodes had bootstrap support >90%. The evolutionary relationships of the populations are as in Papadopulos et al (2021).

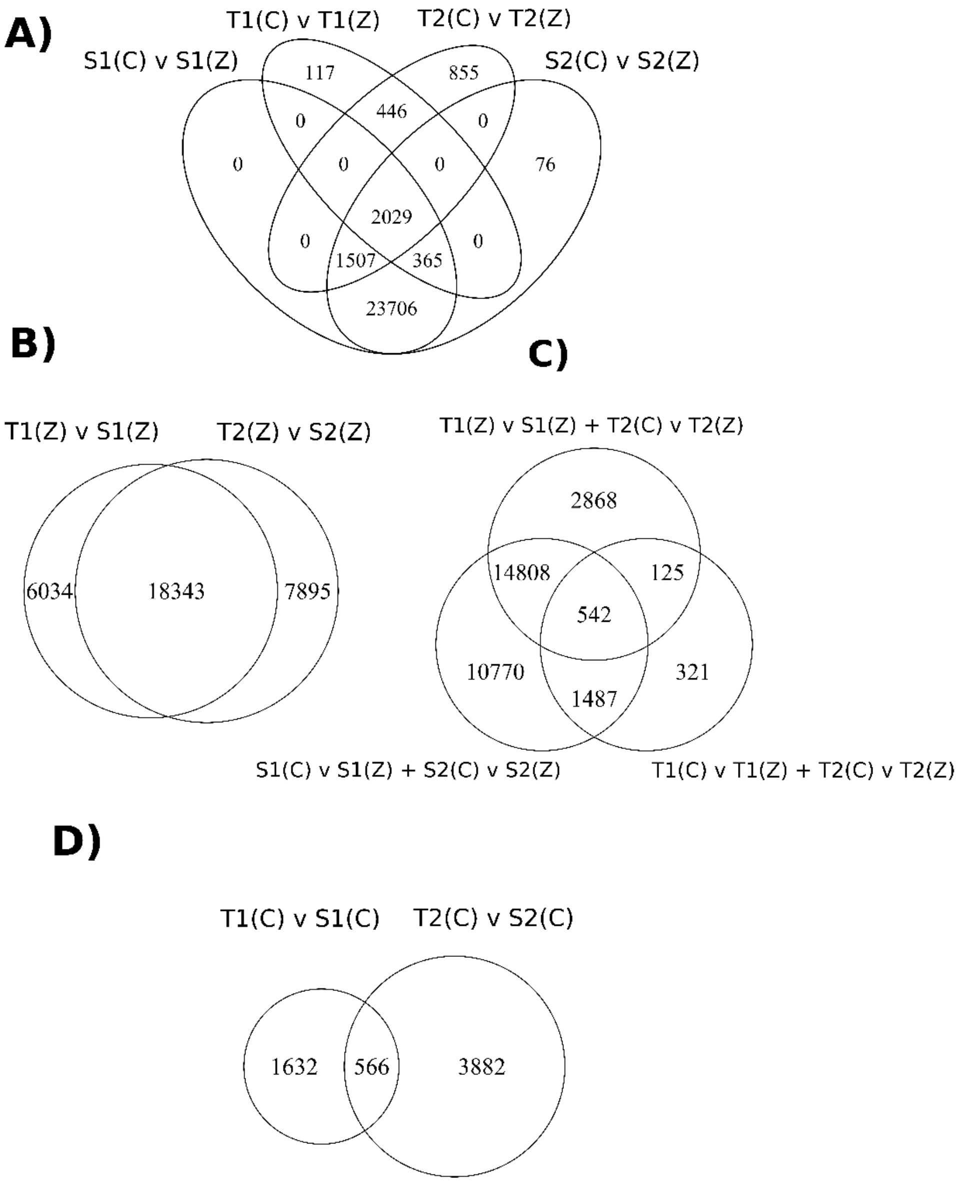
A)
T1(C) v T1(Z)
T2(C) v T2(Z)
S1(C) v S1(Z)
S2(C) v S2(Z)
117
855
0
446
0
0
0
0
76
2029
0
1507
365
0
23706
B)
T1(Z) v S1(Z)
T2(Z) v S2(Z)
6034
18343
7895
C)
T1(Z) v S1(Z) + T2(C) v T2(Z)
2868
14808
125
542
10770
321
1487
S1(C) v S1(Z) + S2(C) v S2(Z)
T1(C) v T1(Z) + T2(C) v T2(Z)
D)
T1(C) v S1(C)
T2(C) v S2(C)
1632
566
3882

**Figure S2: Overlaps between sets of differentially expressed genes.**

Venn diagrams outlining the numbers of significantly differentially expressed genes for a given comparison (separated by a "v"). T1/S1/T2/S2 correspond to populations, (C) corresponds to control conditions, (Z) to zinc conditions: e.g. T1(C) v S1(C) refers to genes differentially expressed between T1 in the control, and S1 in the control. A "+" corresponds to genes that fulfil both sets of criteria: e.g. T1 (C) v T1 (Z) + T2 (C) v T2 (Z) corresponds to the set of genes that are differentially expressed between both i) T1 in the control vs. T1 in the zinc, and ii) T2 in the control vs. T2 in the zinc. Panels illustrate A) Between treatment expression changes within populations, and their overlaps, B) Between ecotype-changes in the zinc in the two geographic groups, and their overlap, C) the overlap between genes differentially expressed in both tolerant populations between conditions, both susceptible populations between conditions, and between both ecotype pairs within the zinc treatment, and D) Between-ecotype changes in the control in the two geographic groups, and their overlap.

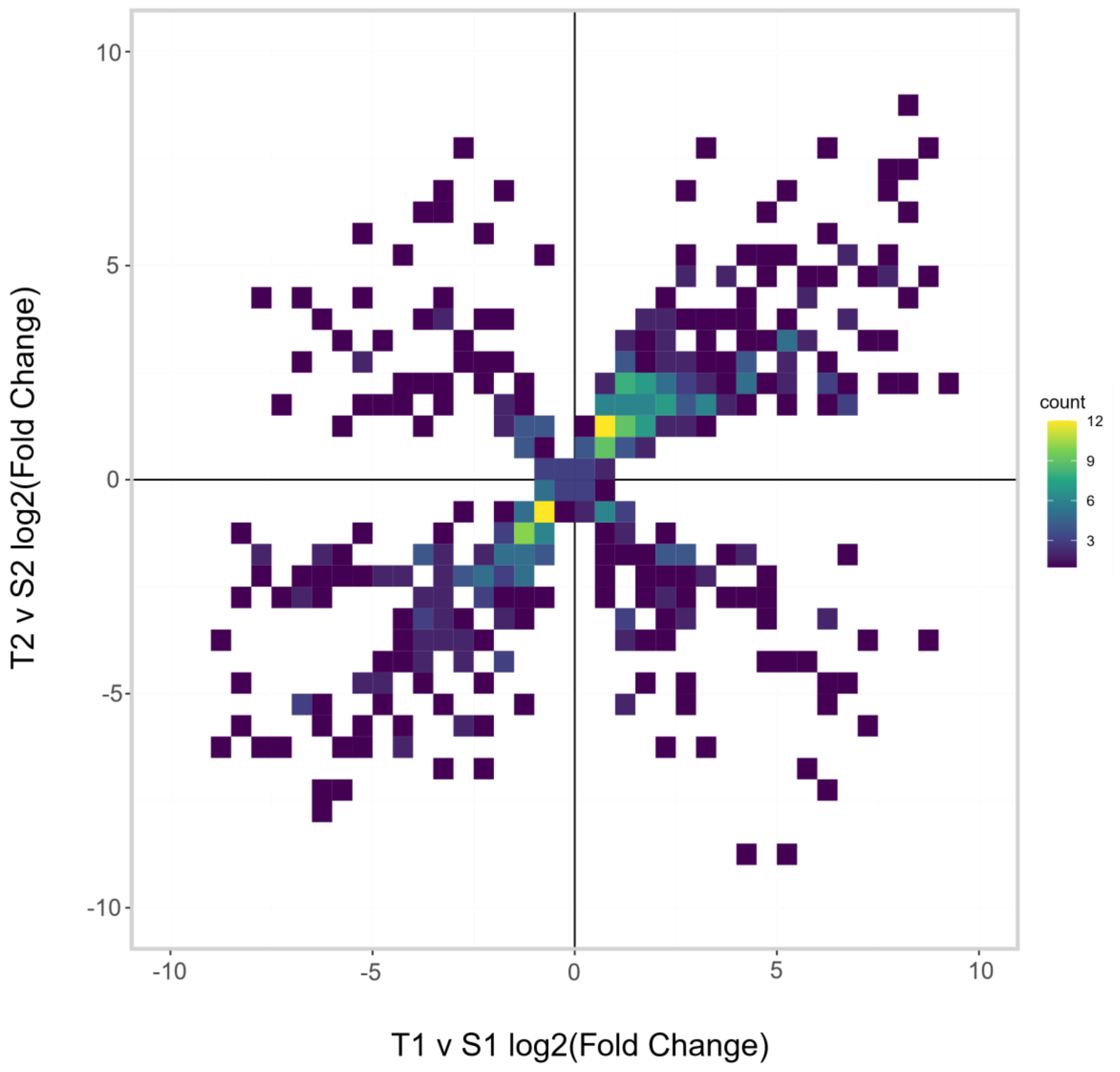


**Figure S3: Fold changes of genes differentially expressed between sensitive and tolerant populations in control conditions.**

Heatmap of log2 transformed shrunken fold changes between T1 and S1 (x axis) and T2 and S2 (y axis) for genes that are significantly differentially expressed between both T1 and S1, and T2 and S2, in the control treatment.

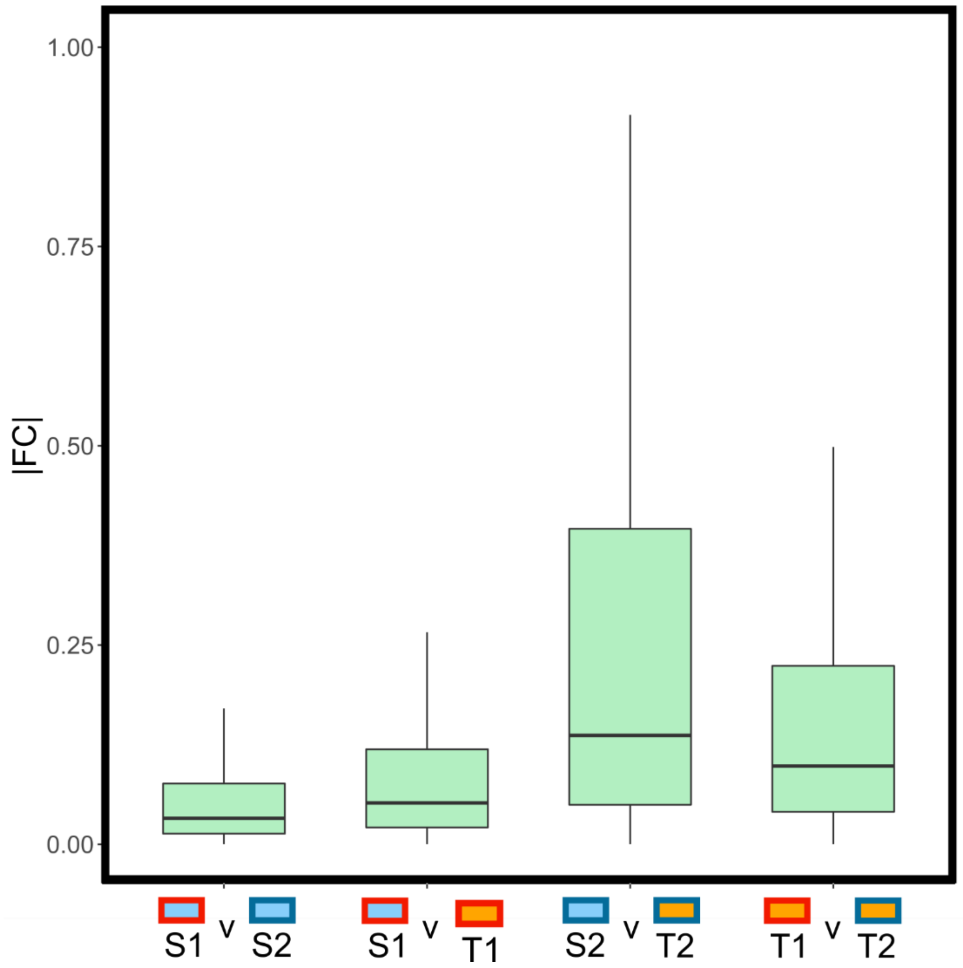


**Figure S4: Overall gene expression differences in the control treatment.**

Boxplots of absolute values of log2 transformed fold changes (|FC|; y axis) between pairs of populations outlined on the x axis, in the control treatment, for all 57,476 genes used in differential expression analysis. Points above 97.25th and below 2.25th percentiles not plotted.

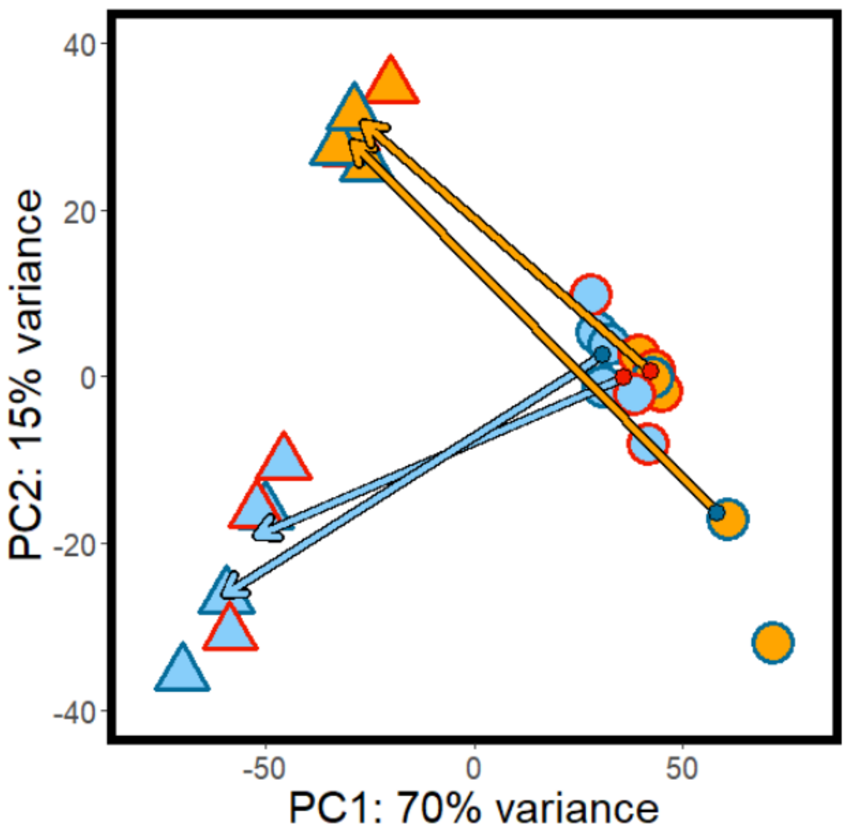


**Figure S5: Principal components analysis of derived plasticity genes.**

Principal components 1 and 2 of variance-transformed counts of all populations in both treatments for genes showing expression differences between treatments in both tolerant populations (i.e. derived plasticity/DP genes). Point fill corresponds to zinc tolerance level (orange = tolerant, blue = sensitive), point border corresponds to geographic pair (red = S1-T1, dark blue = S2-T2). Circles correspond to control treatment, triangles to zinc treatment. Arrows are drawn from the centroid of each population in the control treatment (circle at base of arrow) to the centroid of each population in the zinc treatment (arrowhead). Arrow fill corresponds to zinc tolerance level (orange = tolerant, blue = sensitive) and arrow base point fill corresponds to geographic pair (red = S1-T1, dark blue = S2-T2).

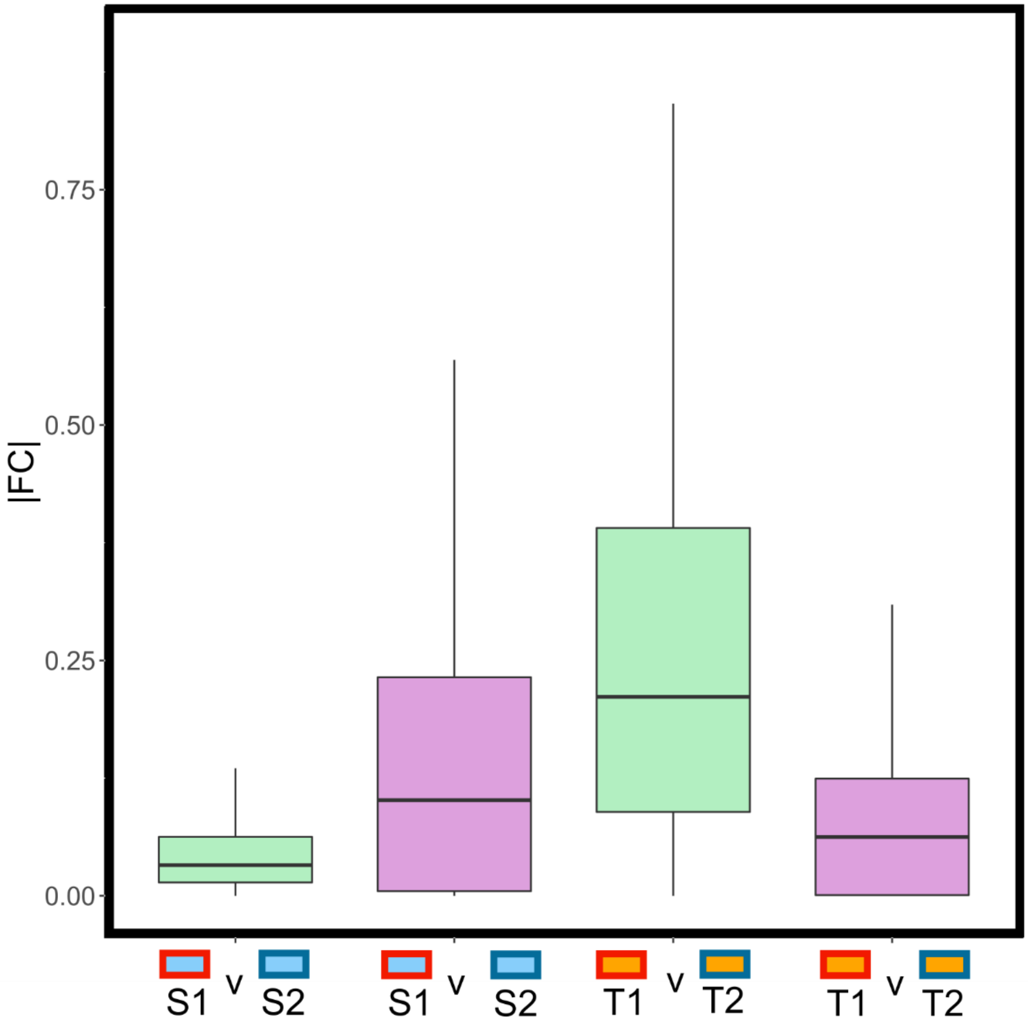


**Figure S6: Gene expression similarity in adaptive plasticity genes across tolerance types and treatments.**

Absolute values of log2 transformed fold changes (|FC|; y axis) between pairs of populations (x axis) for adaptive plasticity genes, either in the control (green) or zinc (purple) conditions. Points above 97.25th and below 2.25th percentiles not plotted.

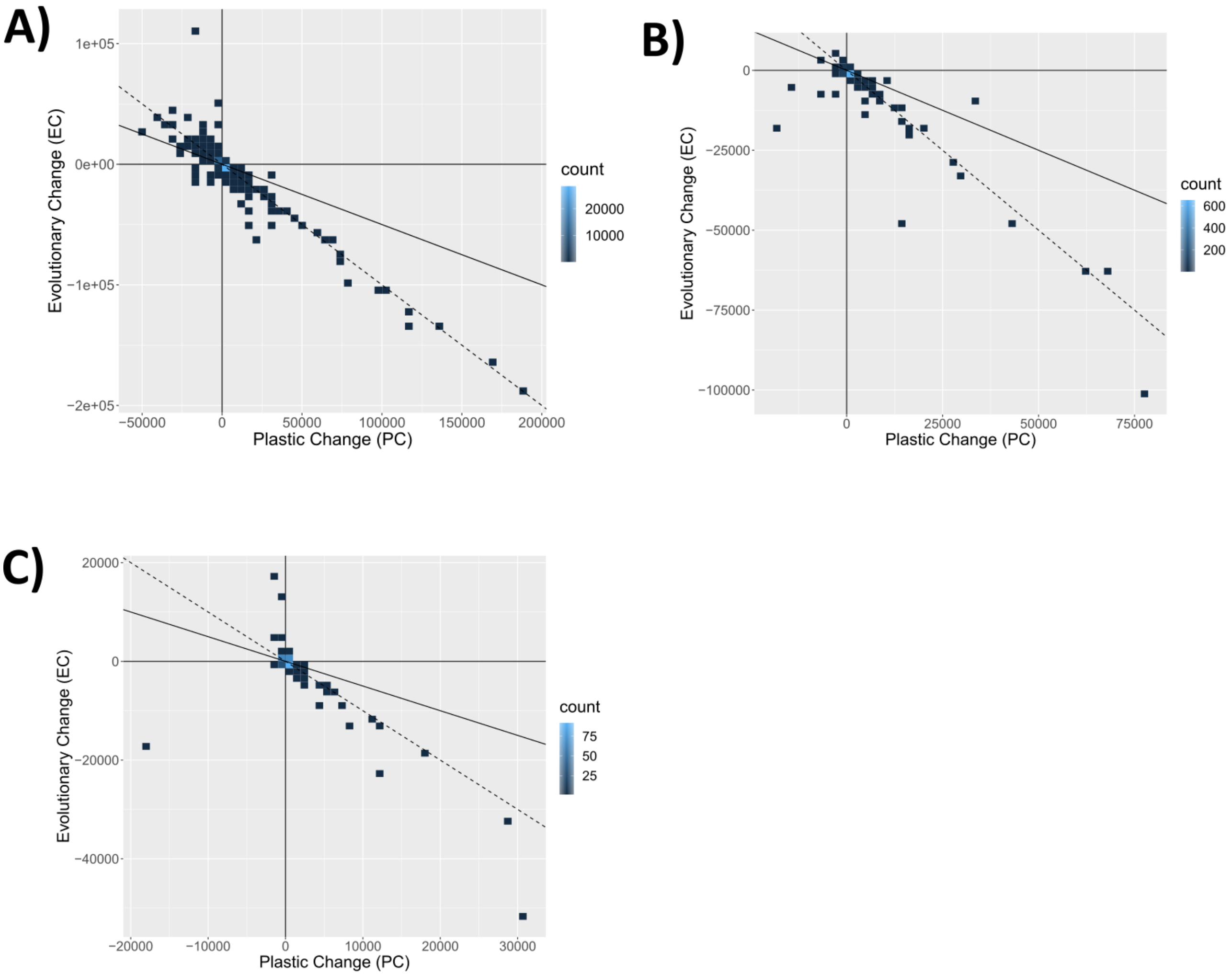


**Figure S7: Relationship between PC and EC for different gene sets.**

Plastic change (PC) vs. evolutionary change (EC) for: A) each gene in the entire transcriptome; B) derived plasticity (DP) genes; and C) genes with constitutive expression differences between ecotypes (CEC).

**Table_S1.tsv: Overrepresented Gene Ontology (GO) terms for genes upregulated in the zinc in S1 and S2.**

For genes upregulated from control to zinc in both S1 and S2 in the same direction, table outlining enriched GO terms. Each row indicates a GO ID with columns indicating: i) the GO_ID; ii) overrepresented p-value; iii) underrepresented p-value; iv) the number of genes in the background set; v) the number of genes in the query set; vi) the GO term; vii) the GO ontology; viii) the over-represented false discovery rate; and ix) the GO term.

**Table_S2.tsv**: **Overrepresented Gene Ontology (GO) terms for genes differentially expressed between S1 and T1, and S2 and T2, in the zinc treatment.**

For genes differentially expressed between T1 and S1, and T2 and S2, in the zinc treatment, in the same direction, table outlining enriched GO terms. Each row indicates a GO ID with columns indicating: i) the GO_ID; ii) overrepresented p-value; iii) underrepresented p-value; iv) the number of genes in the background set; v) the number of genes in the query set; vi) the GO term; vii) the GO ontology; viii) the over represented false discovery rate; and ix) the GO term.

**Table S3: Impact of parametric bootstrapping on response to ancestral plasticity classification.**

Classifications of genes showing substantial ancestral plasticity and evolutionary change from i) the entire transcriptome, ii) genes showing derived plasticity DP genes), iii) genes showing constitutive expression changes (CEC) by their evolutionary response to ancestral plasticity; reversion, overshooting or reinforcement. Listed either without parametric bootstrapping or after parametric bootstrapping (the % of genes followed by the number able to be classified).

| Gene set | Class | No bootstrapping | Bootstrapping |
|---|---|---|---|
| Entire transcriptome | Reversion | 87.8% (38718) | 89.4% (19927) |
| | Overshooting | 5.3% (2339) | 9.3% (2067) |
| | Reinforcement | 6.9% (3055) | 1.3% (293) |
| DP Genes | Reversion | 70.8% (675) | 71.5% (339) |
| | Overshooting | 12.8% (122) | 13.7% (65) |
| | Reinforcement | 16.4% (156) | 14.8% (70) |
| CEC Genes | Reversion | 62.7% (188) | 62.8% (91) |
| | Overshooting | 3.3% (10) | 1.4% (2) |
| | Reinforcement | 34.0% (102) | 35.9% (52) |

**Table S4: Classification of evolutionary response to ancestral plasticity in geographic pairs.**

Table outlining for genes in i) the entire transcriptome, ii) DP genes, iii) genes with derived plasticity in only one of T1 or T2, iii) CEC genes, iv) CEC genes and v) genes with constitutive expression changes in only one of T1 or T2, evolutionary responses to ancestral plasticity. For each gene set, the percentage of genes that could be classified (that showed substantial ancestral plasticity and evolutionary change; see Methods) and the proportions assigned to overshooting, reversions and reinforcement are given. The values are either calculated based on values of $L_o$, $L_p$ and $L_a$ calculated from a) T1 and S1 only (i.e. $L_p$ is average of S1 expression in zinc; $L_a$ is expression of T1 in the zinc etc.), b) T2 and S2 only (i.e. $L_p$ is average of S2 expression in zinc; $L_a$ is expression of T2 in the zinc etc.) and c) the combined values (i.e. $L_p$ is average of S1 and S2 expression in zinc; $L_a$ is expression of T1 and T2 in the zinc etc.; see Methods); note that these are the values used in the text unless otherwise stated.

| | **T1/S1** | **T2/S2** | **Combined** |
|---|---|---|---|
| % of whole transcriptome classified | 75.2% | 75.5% | 76.7% |
| Whole transcriptome classifications:<br>Overshooting<br>Reversion<br>Reinforcement | <br>6.3%<br>85.3%<br>8.4% | <br>5.7%<br>86.6%<br>7.7% | <br>5.3%<br>87.8%<br>6.9% |
| % of DP genes classified | 39.1% | 37.8% | 38.6% |
| DP gene classifications:<br>Overshooting<br>Reversion<br>Reinforcement | <br>11.7%<br>69.8%<br>18.5% | <br>11.8%<br>71.6%<br>16.6% | <br>12.8%<br>70.8%<br>16.4% |
| % of genes with derived plasticity in only one of T1 or T2 classified | 83.5% | 66.1% | NA |
| Genes with derived plasticity in only one of T1 or T2 classifications<br>Overshooting<br>Reversion<br>Reinforcement | <br>4.4%<br>89.9%<br>5.7% | <br>13.1%<br>77.6%<br>9.3% | <br>NA<br>NA<br>NA |
| % of CEC genes classified | 70.5% | 72.6% | 72.6% |
| CEC gene classifications:<br>Overshooting<br>Reversion<br>Reinforcement | <br>4.5%<br>60.1%<br>35.4% | <br>1.7%<br>67%<br>31.3% | <br>3.3%<br>62.7%<br>34.0% |
| % of genes with constitutive expression changes in only one of T1 or T2 classified | 69.0% | 65.1% | NA |
| Genes with constitutive expression changes in only one of T1 or T2 classifications<br>Overshooting<br>Reversion<br>Reinforcement | <br>6.4%<br>77.6%<br>16.0% | <br>7.1%<br>81.8%<br>11.0% | <br>NA<br>NA<br>NA |

**Table_S5.tsv: Overrepresented Gene Ontology (GO) terms for genes significantly upregulated between control and zinc in both T1 and T2.**

For genes upregulated from control to zinc in both S1 and S2 in the same direction, table outlining enriched GO terms. Each row indicates a GO ID with columns indicating: i) the GO_ID; ii) overrepresented p-value; iii) underrepresented p-value; iv) the number of genes in the background set; v) the number of genes in the query set; vi) the GO term; vii) the GO ontology; viii) the over-represented false discovery rate; and ix) the GO term.

**Table_S6.tsv: RNA quantity, quality and reads.**

Table for each RNA sample used: i) sample ID; ii) plant ID; iii) population – plant – treatment code; iv) RNA concentration (Nanodrop ng/µl); v) 260/280 nm absorbance ratio; vi) 260/230 nm absorbance ratio; vii) RNA concentration (Qubit, ng/µl); viii) Elution volume: total (elution 1 + elution 2), µl; ix) Extraction batch (extractions performed in total of 5 batches); x) RNA Integrity Number measured on Aligent Bioanalyser at Beijing Genomics Instititue following shipping; xi) number of raw reads; xii) number of cleaned reads; and xiii) whether the sample was used in transcriptome construction (individual with most total reads per population).

**Table S7: Overlap of adaptive plasticity and constitutive expression differences between geographic pairs.**

For both 1) tolerant vs. susceptible populations in the control (i.e., constitutive expression differences and 2) tolerant populations in the control vs. zinc (i.e., adaptive plasticity genes), the number of genes significantly differentially expressed in: i) geographic pair 1; and ii) geographic pair 2; iii) the overlap between these gene sets; iv) the number of genes in overlapping sets showing expression differences in the same direction; v) the odds ratio; and vi) p-value of this overlap calculated using Fisher's exact test.

| Comparison | Geographic pair 1 | Geographic pair 2 | Overlap | Overlap (same direction) | Odds ratio | p-value |
|---|---|---|---|---|---|---|
| T vs. S (C) | 2198 | 4448 | 566 | 413 | 2.9 | $<2.2 \times 10^{-16}$ |
| T (C vs. Z) | 2957 | 4837 | 2475 | 2473 | 112.3 | $<2.2 \times 10^{-16}$ |

**SI References**

1. A. S. T. Papadopulos, *et al.*, Rapid Parallel Adaptation to Anthropogenic Heavy Metal Pollution. *Mol. Biol. Evol.* (2021).
2. S. A. Andrews, A quality control tool for high throughput sequence data. 2010 (2015).
3. R. K. Patel, M. Jain, NGS QC Toolkit: A Toolkit for Quality Control of Next Generation Sequencing Data. *PLoS One* **7**, e30619 (2012).
4. A. M. Bolger, M. Lohse, B. Usadel, Trimmomatic: A flexible trimmer for Illumina sequence data. *Bioinformatics* **30**, 2114–2120 (2014).
5. B. J. Haas, *et al.*, De novo transcript sequence reconstruction from RNA-seq using the Trinity platform for reference generation and analysis. *Nat. Protoc.* **8**, 1494–1512 (2013).
6. N. L. Bray, H. Pimentel, P. Melsted, L. Pachter, Near-optimal probabilistic RNA-seq quantification. *Nat. Biotechnol.* **34**, 525–527 (2016).
7. M. Krasovec, M. Chester, K. Ridout, D. A. Filatov, The Mutation Rate and the Age of the Sex Chromosomes in Silene latifolia. *Curr. Biol.* **28**, 1832-1838.e4 (2018).
8. W. J. Kent, BLAT---The BLAST-Like Alignment Tool. *Genome Res.* **12**, 656–664 (2002).
9. A. S. T. Papadopulos, M. Chester, K. Ridout, D. A. Filatov, Rapid y degeneration and dosage compensation in plant sex chromosomes. *Proc. Natl. Acad. Sci. U. S.*

*A.* **112**, 13021–13026 (2015).

10. A. Bateman, UniProt: A worldwide hub of protein knowledge. *Nucleic Acids Res.* **47**, D506–D515 (2019).

11. S. F. Altschul, W. Gish, W. Miller, E. W. Myers, D. J. Lipman, Basic local alignment search tool. *J. Mol. Biol.* **215**, 403–410 (1990).

12. M. Seppey, M. Manni, E. M. Zdobnov, "BUSCO: Assessing genome assembly and annotation completeness" in *Methods in Molecular Biology*, (Humana Press Inc., 2019), pp. 227–245.

13. R. D. Finn, *et al.*, Pfam: The protein families database. *Nucleic Acids Res.* **42**, D222 (2014).

14. J. Mistry, R. D. Finn, S. R. Eddy, A. Bateman, M. Punta, Challenges in homology search: HMMER3 and convergent evolution of coiled-coil regions. *Nucleic Acids Res.* **41**, e121–e121 (2013).

15. D. M. Bryant, *et al.*, A Tissue-Mapped Axolotl De Novo Transcriptome Enables Identification of Limb Regeneration Factors. *Cell Rep.* **18**, 762–776 (2017).

16. R Core Team, R: A language and environment for statistical computing (2013).

17. C. Soneson, M. I. Love, M. D. Robinson, Differential analyses for RNA-seq: transcript-level estimates improve gene-level inferences. *F1000Research* **4**, 1521 (2015).

18. M. I. Love, W. Huber, S. Anders, Moderated estimation of fold change and dispersion for RNA-seq data with DESeq2. *Genome Biol.* **15**, 550 (2014).

19. M. Stephens, False discovery rates: A new deal. *Biostatistics* **18**, 275–294 (2017).

20. M. D. Young, M. J. Wakefield, G. K. Smyth, A. Oshlack, Gene ontology analysis

for RNA-seq: accounting for selection bias. *Genome Biol.* **11**, R14 (2010).

21. C. K. Ghalambor, *et al.*, Non-adaptive plasticity potentiates rapid adaptive evolution of gene expression in nature. *Nature* **525**, 372–375 (2015).
22. W. C. Ho, J. Zhang, Evolutionary adaptations to new environments generally reverse plastic phenotypic changes. *Nat. Commun.* **9**, 1–11 (2018).
23. W. C. Ho, J. Zhang, Genetic Gene Expression Changes during Environmental Adaptations Tend to Reverse Plastic Changes Even after the Correction for Statistical Nonindependence. *Mol. Biol. Evol.* **36**, 604–612 (2019).
24. D. Kim, J. M. Paggi, C. Park, C. Bennett, S. L. Salzberg, Graph-based genome alignment and genotyping with HISAT2 and HISAT-genotype. *Nat. Biotechnol.* **37**, 907–915 (2019).
25. H. Li, *et al.*, The Sequence Alignment/Map format and SAMtools. *Bioinformatics* **25**, 2078–2079 (2009).
26. T. H. Lee, H. Guo, X. Wang, C. Kim, A. H. Paterson, SNPhylo: A pipeline to construct a phylogenetic tree from huge SNP data. *BMC Genomics* **15**, 162 (2014).
